# Mechanosensitive FHL2 tunes endothelial function

**DOI:** 10.1101/2024.06.16.599227

**Authors:** Shailaja Seetharaman, John Devany, Ha Ram Kim, Emma van Bodegraven, Theresa Chmiel, Shentu Tzu-Pin, Wen-hung Chou, Yun Fang, Margaret Lise Gardel

## Abstract

Endothelial tissues are essential mechanosensors in the vasculature and facilitate adaptation to various blood flow-induced mechanical cues. Defects in endothelial mechanoresponses can perturb tissue remodelling and functions leading to cardiovascular disease progression. In this context, the precise mechanisms of endothelial mechanoresponses contributing to normal and diseased tissue functioning remain elusive. Here, we sought to uncover how flow-mediated transcriptional regulation drives endothelial mechanoresponses in healthy and atherosclerotic-prone tissues. Using bulk RNA sequencing, we identify novel mechanosensitive genes in response to healthy unidirectional flow (UF) and athero-prone disturbed flow (DF). We find that the transcription as well as protein expression of Four-and-a-half LIM protein 2 (FHL2) are enriched in athero-prone DF both *in vitro* and *in vivo*. We then demonstrate that the exogenous expression of FHL2 is necessary and sufficient to drive discontinuous adherens junction morphology and increased tissue permeability. This athero-prone phenotype requires the force-sensitive binding of FHL2 to actin. In turn, the force-dependent localisation of FHL2 to stress fibres promotes microtubule dynamics to release the RhoGEF, GEF-H1, and activate the Rho-ROCK pathway. Thus, we unravelled a novel mechanochemical feedback wherein force-dependent FHL2 localisation promotes hypercontractility. This misregulated mechanoresponse creates highly permeable tissues, depicting classic hallmarks of atherosclerosis progression. Overall, we highlight crucial functions for the FHL2 force-sensitivity in tuning multi-scale endothelial mechanoresponses.

## Introduction

Endothelial cells in the vasculature constantly sense and respond to mechanical cues, including shear stress caused by blood flow and stretch during vascular dilation^1,2^. Such mechanoresponses are crucial for the formation and maintenance of blood vessels during development, angiogenesis, and homeostasis^3,4^. However, mechanical cues within the vasculature also drive the onset and progression of cardiovascular diseases including atherosclerosis, stenosis, and aneurysms^5^. For instance, changes in blood flow profiles mimicking atherosclerosis promote drastic phenotypical remodelling of the endothelium and subsequently, increase tissue permeability and inflammation^6^. Concurrently, changes in transcriptional profiles further promote aberrant phenotypes through subcellular signalling that alters cell morphology and tissue permeability^7^. These phenotypes and gene expression patterns are key hallmarks of endothelial dysfunction^8,9^. Thus, endothelial physiology requires multiscale mechanobiological processes controlling cell morphology, adhesion, and transcriptional state. Owing to these complexities, there exist very few therapies for cardiovascular diseases that directly target vascular cell dysfunction^10^. Therefore, we sought to understand how endothelial mechanoresponses shape the onset and progression of atherosclerotic phenotypes.

Endothelial mechanotransduction requires specialised cell surface receptors (e.g., membrane receptors, cilia, and ion channels^11–16^), but also mechanical contacts at adhesions to the extracellular matrix (focal adhesions) or neighbouring cells (adherens junctions) and throughout the cytoskeleton^17–20^. These mechanosensitive processes are crucial for endothelial tissue phenotypes and functions, including changes in adhesion morphology and permeability^21–23^. In healthy endothelial tissues, permeability is controlled by forming and maintaining stable adherens junctions^24,25^. However, the activation of the RhoGTPase signalling pathway, via RhoA, enhances actomyosin contractility, and such hypercontractility is often seen in atherosclerotic tissues^26^. This, in turn, generates discontinuous adherens junctions, enhanced tissue permeability, and induction of classic biochemical and transcriptional signatures of endothelial tissue dysfunction^27–29^. However, the mechanisms by which subcellular mechanotransduction at actomyosin and adhesions alter endothelial tissue function remain unclear. Furthermore, how mechanotransduction pathways, that can operate both through rapid biochemical signals and long-term transcriptional regulation, control cell adhesion and morphology remain elusive. Such multi-scale integration of mechanotransduction in the endothelium is critical for advancing our capabilities to precisely engineer tissue-scale physiology.

Here, we identify Four-and-a-half LIM domain protein 2 (FHL2) as a key regulator of multiscale mechanotransduction that drives the onset of athero-prone endothelial phenotypes. Using RNA sequencing, we identify mechanosensitive genes in human aortic endothelial tissues subjected to flow profiles mimicking athero-protective unidirectional flow (UF) and athero-prone disturbed flow (DF). We observed differential expression of numerous adhesion- and actomyosin-related genes, with the LIM (<u>L</u>in11, Isl-1 & <u>M</u>ec-3) domain family of proteins being the top enriched gene family within these categories. Using human aortic endothelial cells and mouse models, we find that *FHL2* is transcriptionally upregulated in athero-prone flow conditions. We demonstrate that FHL2 expression is sufficient to drive hallmarks of an athero-prone phenotype, including discontinuous adherens junctions and enhanced tissue permeability. We show that increased FHL2 expression results in hypercontractility, as seen in athero-prone conditions. We demonstrate that FHL2 tunes contractility through crosstalk between the actin and microtubule cytoskeleton and requires force-activated binding of FHL2 to actin. Specifically, FHL2 alters microtubule dynamics and organisation to promote contractility via the release of a Rho guanine nucleotide exchange factor, GEF-H1 from microtubules into the cytosol, where it activates RhoA. Subsequent FHL2-mediated hypercontractility, via reinforcement of a positive feedback, yields athero-prone phenotypes. Broadly, our results uncover a novel mechanochemical feedback providing crucial mechanistic insight into how endothelial mechanoresponses go awry to promote atherosclerotic phenotypes.

## Results

### Transcriptomics reveals novel mechanosensitive genes in endothelial tissue response to flow

In healthy arteries resistant to atherosclerosis, blood flow is laminar and unidirectional; however, in arterial sites prone to atherosclerosis such as bifurcations, branches, and curvatures^30,31^, vascular endothelial cells are constantly activated by disturbed blood flow. An example of this is the human carotid artery bifurcation, which most commonly develops atherosclerotic plaques^32,33^ (Fig. 1A). Within this site, recent work has identified two arterial waveforms: “athero-protective” and “athero-prone” flows, which represent blood flow in the distal internal carotid artery and carotid sinus respectively (Fig. 1A)^33^. These hemodynamic waveforms were captured *in vitro* using a motorised cone-plate system^33–35^. Specifically, unidirectional flow (UF) with a high shear stress (average of 20 dyn/cm^2^, with a peak of ∼40 dyn/cm^2^) mimics the athero-protective hemodynamic force in the distal carotid artery, and an oscillatory disturbed flow (DF) with a low shear stress (average of 2 dyn/cm^2^, with a peak of ∼10 dyn/cm^2^) represents flows in athero-prone arterial regions (Fig. 1A, 1B)^33^.

**Figure 1:**
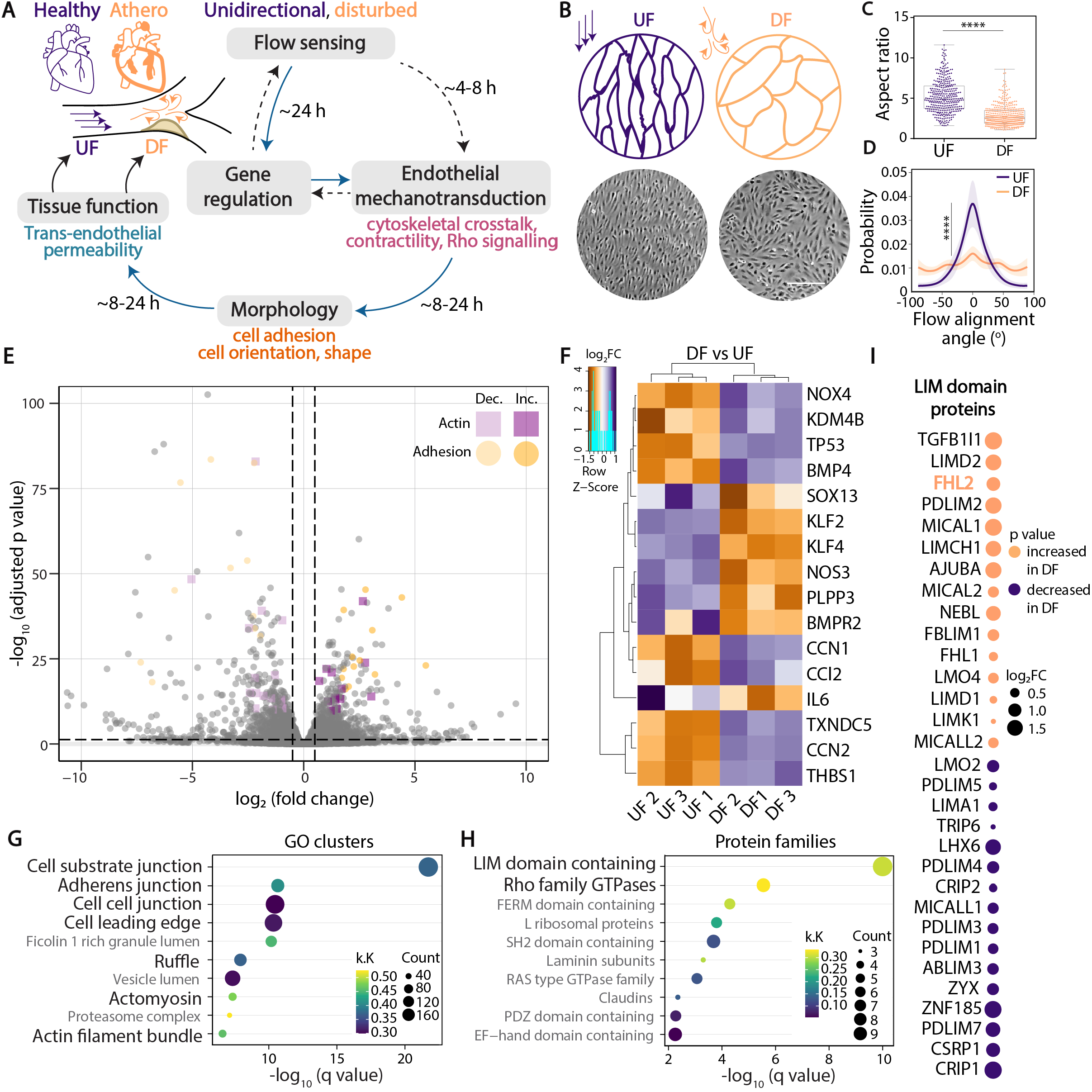
Identification of flow-sensitive gene expression profiles in endothelial cells. **(A)** Schematic showing the interplay between gene expression and endothelial mechanotransduction that facilitates multi-scale tissue adaptation, morphology, and function. In particular, hypercontractility of tissues through RhoGTPase mechano-signalling, cell-cell adhesion formation, and gene expression, collectively orchestrate tissue functions including permeability. **(B)** TeloHAECs (human aortic endothelial cells) subjected to two flow profiles: healthy laminar unidirectional flow (UF) and athero-prone disturbed flow (DF). Brightfield images of cells under UF and DF are shown. **(C)** Graph shows the aspect ratio (ratio of the major to the minor axis of a cell) of cells under UF and DF. In the box-and-whisker plot, the box extends from the 25th to the 75th percentile, the whiskers show the minimum and maximum values, and the line within the box represents the median. N=3 independent experiments (n=361 for UF, and n=425 for DF). **(D)** From brightfield images, the alignment of cells under UF and DF are calculated by local gradient orientation, and represented as the probability distribution of cells with the flow alignment angles (centred around 0°). Shaded error bars representing standard deviation are shown. N=3 independent experiments (n=30 for UF, and n=32 for DF; fields of view, with >100 cells/field). **(E)** Volcano plot shows differentially expressed genes (DEGs) between TeloHAECs under UF and DF (dotted lines indicate log_2_FC 0.5 and -log_10_(adjusted p-value = 0.05)). 1260/12539 genes were significantly upregulated in DF (adjusted p-value < 0.05, log_2_FC > 0.5), and 2079/12539 genes were significantly downregulated in DF (adjusted p-value < 0.05, log_2_FC < 0.5). Genes annotated to adhesion-related GO terms (Supplementary Table 2; dark orange circles: logFC > 0, light orange circles: logFC < 0) and actin-related Gene Ontology (GO) terms (Supplementary Table 2; dark purple squares: logFC > 0, light purple squares: logFC < 0). **(F)** Heatmap showing Z-scored rlog transformed count data (log_2_ transformation and normalization with respect to library size) normalized of flow-sensitive genes across samples. **(G)** Top 10 most significantly enriched cellular compartment GO terms. Dot size = gene set size (count = number of genes; k.K. = number of DEG genes in GO term/number of genes in the GO term, qvalue = Benjamini-Hochberg corrected p-value (multiple hypothesis test correction). **(H)** All DEGs which were present within adhesion- and actin-related GO terms were tested for overrepresentation in gene families. The top 10 most significantly enriched gene families are shown. **(I)** Dotplot showing all genes significantly different between TeloHAECs under UF and DF which encode for LIM domain proteins ordered by adjusted p-value and logFC. The size of the dots indicates absolute logFC values. Four-and-a-half LIM protein 2 (FHL2) is highlighted in orange as an upregulated gene in cells under DF. For E-F, N=3 independent experiments. **Statistical tests:** (C) unpaired Student’s t-test followed by Mann-Whitney test (two-tailed). (D) Kolmogorov-Smirnov test. ****p<0.0001.

Using this model, we applied UF and DF for 24 h to a monolayer of human aortic endothelial cells (TeloHAEC) to characterise the tissue-scale response (Fig. 1B). Consistent with previous studies, we observed that under UF, endothelial cells elongated and aligned in the direction of flow (Fig. 1B, 1C, 1D). In contrast, cell elongation and tissue alignment were lost in cells subjected to DF (Fig. 1B, 1C, 1D). These morphological characteristics are consistent with the phenotypes seen *in vivo* of the descending thoracic aorta and the aortic arch (inner curvature) that experience UF and DF respectively^36^.

A limited number of transcriptomic studies have been carried out in endothelial cells experiencing static, low or high shear stress^33,37,38^. Building on a previous study^39^, the flow profiles used here provide a reliable comparison between two distinct conditions of healthy UF and athero-prone DF flows. Using this approach, we performed bulk RNA sequencing to identify the transcriptional changes accompanying endothelial mechanoresponses to 24 h of flow. This transcriptomic screen revealed approximately 3339 differentially expressed genes (DEGs; adjusted p-value <0.05), with 1260 and 2079 genes upregulated in DF and UF respectively (Fig. 1E; Table S1). Under these conditions, we depict known endothelial flow-mediated transcriptional changes, including Krüppel-like factors *KLF2* and *KLF4*, and nitric oxide synthase 3 (*NOS3*) (Fig 1F)^10,40^.

Further, gene ontology (GO) analysis on all DEGs indicated that the most enriched gene sets in the “cellular component CC” category were related to “actin” and “adhesion” (Fig. 1E, 1G; Table S2; actin- and adhesion-related gene sets are defined in the methods). The top DEGs belonging to these two categories are listed in Fig. S1A, S1B. Amongst the DEGs annotated to the “actin” and “adhesion” terms, we observed that LIM domain-containing proteins and RhoGTPases were the top enriched gene families (Fig. 1H). In fact, we found that 31 genes in the LIM domain containing family regulated in response to changes in flow profiles (Fig. 1I, S1C; arranged based on adjusted p-values). As LIM domain and RhoGTPase families of proteins are crucial regulators of mechanotransduction in adherent cells^41^, we chose to focus on the role of these genes in the context of flow-induced endothelial dysfunction.

### Disturbed flow induces FHL2 endothelial expression *in vitro* and *in vivo*

The LIM domain protein family contains ∼70 genes divided into 14 classes, which play critical roles in cytoskeletal organisation, adhesion, and migration, and broadly contribute to diseases including cancer. In numerous cases, the LIM domain proteins exhibit force-dependent localisation to adherens junctions, focal adhesions, and the actin cytoskeleton and have been implicated in mechanotransduction at these structures^42–50^. However, only a limited number of LIM-mediated mechanotransduction pathways are known. Some LIM domain proteins are thought to act as transcriptional co-factors^51^; however, the mechanisms by which their transcriptional activity tunes mechanotransduction have not been explored. Thus, the functions of mechanosensitive LIM proteins and LIM-dependent downstream signalling cascades remain largely elusive, particularly in the case of cardiovascular systems. Interestingly, one of the most upregulated genes in our screen was FHL2, which is a LIM-only protein (Fig. 1I). FHL2 is both highly expressed in the heart^52^ and exhibits force-sensitive recruitment to actin^46^. Further, studies have shown that mice with Fhl2 deletion developed smaller atherosclerotic plaques and exhibited lower inflammation^53–56^, suggesting a role for FHL2 in atherosclerotic disease progression.

To assess the regulation of Fhl2 *in vivo*, we employed a partial carotid artery ligation mouse model to induce endothelial dysfunction. Here, partial ligation of the left carotid artery induces acute disturbed flow and low shear stress in the artery, resulting in the formation of an atherosclerotic plaque in approximately 2 weeks post-surgery^57,58^ (Fig. 2A). Most of the effects seen in arteries at 24-48 h post-ligation are attributed to the DF alone, and not atherosclerosis-induced endothelial dysfunction. Exploiting this powerful model system, we isolated the endothelium-enriched intima of contralateral (non-ligated right carotid artery, as control) and ligated mouse arteries 48 h post-surgery, and subsequently extracted RNA from these mouse intima tissues. Quantitative PCR revealed that the levels of crucial transcription factors and well-known flow-sensitive genes, *Klf2* and *Nos3*, are downregulated by ∼3-4 fold in the ligated arteries (Fig. 2B), which is consistent with Fig. 1 and previous work^10^. In ligated arteries with DF, we observed that *Fhl2* expression increased by >3-fold compared to the contralateral arteries (Fig. 2B). Next, we tested whether these transcriptional changes correlated with protein expression variations in mouse arteries. By immunofluorescence staining for FHL2 in partial ligation mouse models, we observed that the endothelial lining of the ligated mouse artery had a striking increase in FHL2 expression compared to the contralateral control arteries (Fig. 2C). This strongly suggests that endothelial FHL2 expression precedes the onset of an atherosclerotic plaque.

**Figure 2:**
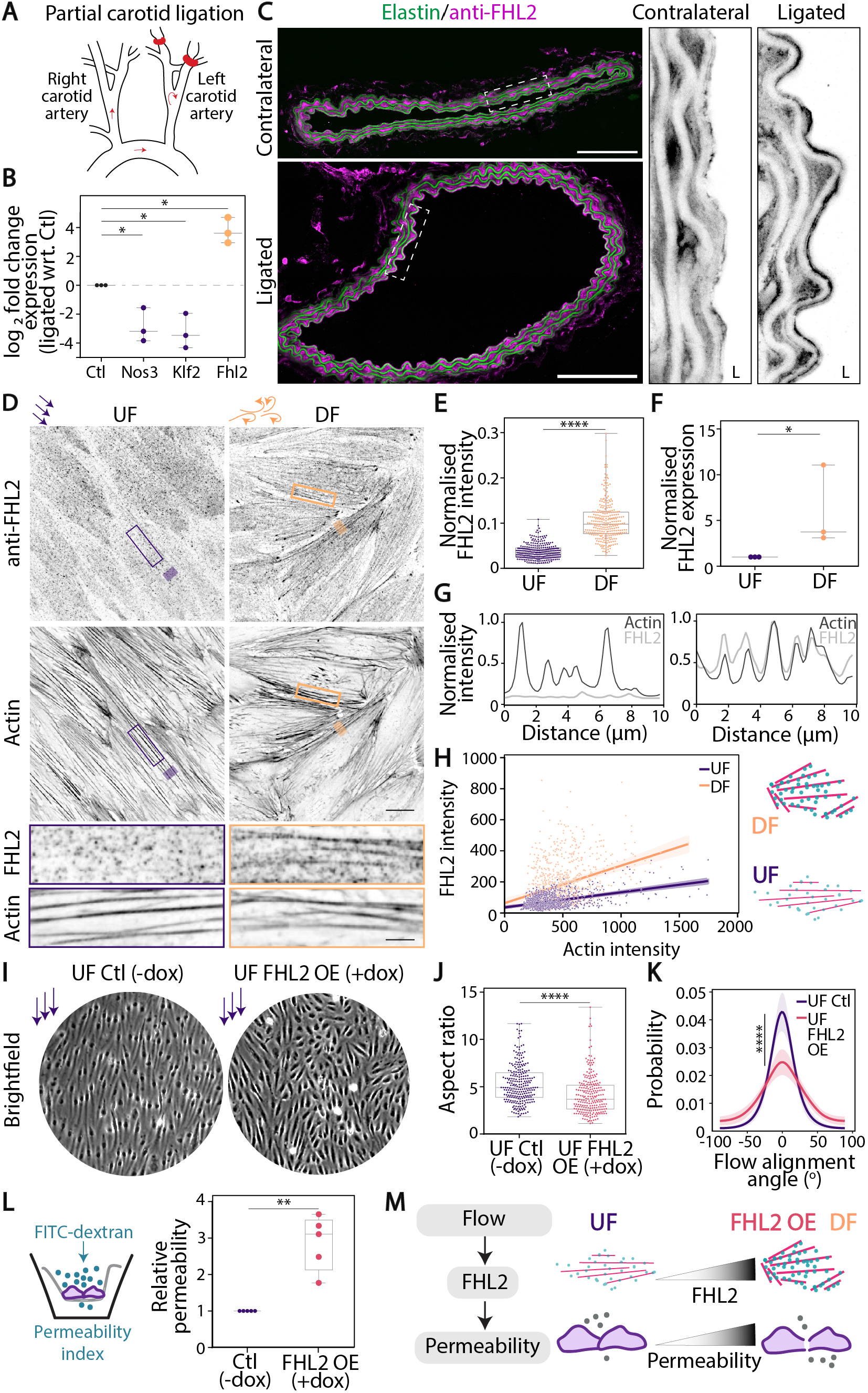
FHL2 expression and localisation to stress fibres is enriched in cells under athero-prone DF. **(A)** Partial ligation of the left carotid artery of the aortic arch in mice (indicated with a red band). The right carotid artery is labelled. 48 h post ligation surgery, mice were euthanised and RNA was extracted from the intima of both carotid arteries (with the contralateral artery as control). **(B)** Graph shows the fold change expression of genes with respect to the expression in control (Ctl) arteries, as measured by qPCR. Two flow-sensitive genes nitric oxide synthase 3 (*Nos3*) and Krüppel-like Factor 2 (*Klf2*) are shown as positive controls/validations. The fold change in *Fhl2* expression is depicted. N=3 mice. **(C)** Confocal images of cross sections of contralateral and ligated mouse arteries stained for FHL2, imaged along with Elastin. Zoomed-in images of FHL2 staining from each arterial section are shown, with ‘L’ indicating the lumen and endothelial side. N=2 mice (n>6 sections per condition). **(D)** Inverted contrast confocal images of TeloHAECs under UF and DF, stained for actin to label stress fibres and FHL2. The regions marked in purple and orange boxes are zoomed in below. **(E)** Normalised expression of FHL2, as measured by qPCR, in TeloHAECs under DF compared to cells under UF. N=3 independent experiments. **(F)** Fluorescence intensity of FHL2 normalised to actin intensity of each cell, in TeloHAECs under UF and DF. N=2 independent experiments (n=290 for UF, 289 for DF). **(G)** Line scans showing FHL2 and actin intensities from a representative region of a stress fibre. **(H)** Graph showing actin intensity vs. FHL2 intensity in a region of a stress fibre. N=3 independent experiments (n=1026 SF regions for UF, 917 SF regions for DF). Slopes are 0.09 and 0.24 for UF and DF respectively. Shaded error bars are shown. **(I)** Brightfield images of Control (Ctl; -dox) and FHL2 overexpressing (OE; +dox) TeloHAECs under UF are shown. **(J)** Graph shows the aspect ratio (ratio of the major to the minor axis of a cell) of control and FHL2 OE (+dox) cells under UF. N=2 independent experiments (n=253 for Ctl, 236 for FHL2 OE). **(K)** From brightfield images, the alignment of control and FHL2 OE (+dox) cells under UF are calculated by local gradient orientation, and represented as the probability distribution of cells with the flow alignment angles (centred around 0°). Shaded error bars representing standard deviation are shown. N=3 independent experiments (n=22 for UF Ctl, and n=26 for UF FHL2 OE; fields of view, with >100 cells/field). **(L)** Schematic of a transwell permeability assay, where dextran-FITC (green) was added to the top chamber, and permeability was calculated by measuring green fluorescence in the collected media from the top and bottom chambers. Graph shows relative permeability levels of FHL2 OE cells, normalised with respect to the Ctl. N=5 independent experiments. **(M)** Schematic showing that FHL2 levels increase in cells under DF. With increased FHL2 expression, endothelial tissues exhibit high permeability and the lack of cell alignment with flow (disrupted tissue phenotypes), while cells with lower levels of FHL2 (UF or Ctl) have normal physiological tissue phenotypes. In the box-and-whisker plots, the box extends from the 25th to the 75th percentile, the whiskers show the minimum and maximum values, and the line within the box represents the median. **Scale:** (B) 100 µm, (D) 20 µm; inset 5 µm. **Statistical tests:** (A) One-way ANOVA followed by Tukey’s multiple comparison’s test. (E, F, K) Student’s t-test followed by Mann-Whitney test. (L) Kolmogorov-Smirnov test. (M) Paired t-test. *p<0.05, **p<0.005 ****p<0.0001.

Next, we investigated whether our findings from mouse tissues were observable in monolayers of endothelial cells subjected to 24 hrs of UF and DF profiles, representing athero-protective and athero-prone conditions (as described in Fig. 1). High resolution confocal imaging of FHL2 and actin allowed for quantification of changes in overall FHL2 protein levels and its subcellular localization (Fig. 2D). FHL2 protein levels averaged across a cell increased by two-fold in DF (Fig. 2E). This was also validated via western blotting (Fig. S2A, S2B). Consistent with RNA seq data and in vivo mouse intima, qPCR revealed that FHL2 transcription increased by ∼3-fold in DF (Fig. 2F). Concurrently, the subcellular distribution of FHL2 also changed in response to flow. To quantify this, intensity profiles across stress fibres in each condition were obtained (Fig. 2D, 2G). While robust actin stress fibres were seen in both UF and DF conditions (Fig. 2D), stress fibres in DF conditions were strongly enriched FHL2 (Fig. 2D, 2G). To explore whether changes in FHL2 localization arose from differences in stress fibre actin density, we plotted FHL2 intensity as a function of actin intensity for a variety of stress fibres. We observed large stress fibres with high FHL2 intensity in DF cells. On the contrary, in UF, majority of stress fibres were smaller, and the occasional large stress fibres did not contain a proportionately high FHL2 intensity as was seen in DF cells. Taken together with previous work showing FHL2 binds tensed actin^46^, our results suggest that in athero-prone DF, actomyosin bundles are highly tense and facilitate the binding of FHL2 to stress fibres (Fig. 2D, 2H).

We then investigated whether the changes in FHL2 expression are sufficient to drive endothelial dysfunction, even under athero-protective flow profiles. We generated a doxycycline (dox)- inducible endothelial cell line overexpressing FHL2 (TeloHAEC FHL2-mEmerald; FHL2 OE), where the FHL2 protein levels measured by western blotting (Fig. S2C, S2D) or immunofluorescence (Fig. S2E, S2F) were approximately two times higher than in control cells. This difference in FHL2 expression is similar to the fold change between endogenous FHL2 levels in cells under UF and DF (Fig. 2E, 2F). We also confirmed that the localisation of FHL2 in the overexpression cells resembled the endogenous protein (Fig. S2E; movie 1). We observed that FHL2-overexpressing cells subjected to UF were less elongated compared to the control cells (Fig. 2I, 2J). Furthermore, FHL2 overexpression reduced cell alignment in the direction of flow (Fig. 2K). Thus, FHL2 upregulation is sufficient to drive the cell morphological phenotypes in athero-prone DF, even when subjected to athero-protective UF.

A well-established readout of endothelial dysfunction in *in vivo* as well as *in vitro* athero-prone DF models is tissue permeability or leakiness^59,60^. Due to the FHL2-mediated perturbations in cell morphology and alignment, we specifically focused on the role of FHL2 in tissue-scale endothelial function. Using transwell assays to assess the amount of fluorescent dextran that crossed the endothelial monolayer, we observed that *in vitro* endothelial tissues with FHL2 overexpression were ∼3 times more permeable compared to the control endothelial monolayers (Fig. 2L). Taken together, our results from *in vitro* and *in vivo* experiments strongly demonstrate that FHL2 is upregulated in response to athero-prone DF. We further show that increased FHL2 expression, as seen in DF, promotes tissue permeability (Fig. 2M). This suggests a crucial role for FHL2 transcriptional regulation and localisation in regulating endothelial mechanoresponses to DF at the onset of atherosclerosis.

### FHL2 expression drives the formation of discontinuous focal adherens junctions

Endothelial permeability and barrier function are tightly regulated by the stability and integrity of adherens junctions^22,25,61^. In atherosclerotic arteries, the breakdown of cell junctions is a classical marker of endothelial dysfunction^10,33^. By visualising VE-Cadherin in endothelial cells under UF and DF, three junction morphologies are observed (Fig. 3A): 1) linear junctions which are thin, intact, well-formed and stable; 2) focal adherens junctions which represent the dynamic and discontinuous junctions, with parts of the junctions along cell borders being broken^62,63^; and 3) reticular junctions which are wider, overlapping and honeycomb-like^64^.

**Figure 3:**
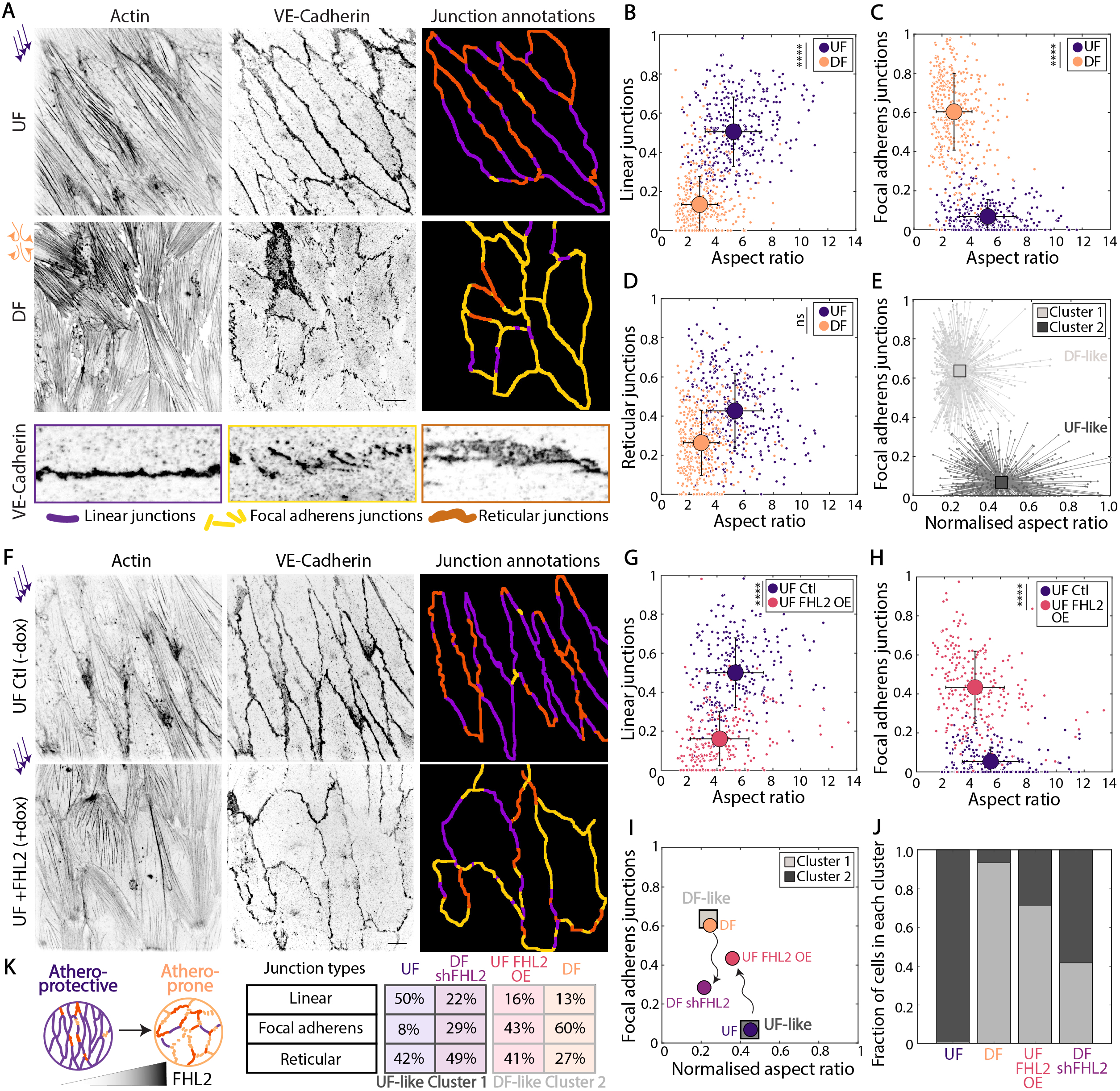
FHL2 overexpression drives focal adherens junction formation, similar to DF cell clusters. **(A)** Inverted contrast confocal images of TeloHAECs under UF and DF, stained for actin and vascular endothelial cadherin, VE-Cadherin (to label adherens junctions). Three cell junction types were annotated using VE-Cadherin staining: i. linear junctions, ii. reticular junctions, iii. focal adherens junctions. Examples of each junction type are shown as zoomed-in inverted contrast images on the bottom. **(B, C, D)** Graphs showing the aspect ratio (ratio of the major to the minor axis of a cell) vs. the fraction of the cell perimeter occupied by linear junctions (B), focal adherens junctions (C), reticular junctions (D). N=3 independent experiments (n=356 for UF, 417 for DF). **(E)** Using measurements of aspect ratio and fractions of the three junction types in B-D, cells were clustered using k-means, with k=2. Two clusters and the corresponding data points are shown. N>3 independent experiments (n=1179 cells clustered into cluster 1 or 2). **(F)** Inverted contrast confocal images of control (Ctl; -dox) and FHL2-overexpressing (OE; +dox) TeloHAECs under UF, stained for phalloidin and VE-Cadherin. Annotated junctions are depicted as in (A). **(G, H, I)** Graphs showing the aspect ratio (ratio of the major to the minor axis of a cell) vs. the fraction of the cell perimeter occupied by linear junctions (B), focal adherens junctions (C), reticular junctions (D). N=2 independent experiments (n=253 for UF Ctl, 256 for UF FHL2 OE). The UF Ctl means were pooled from 5 experiments for statistical tests. **(J)** Centroids of clusters 1 and 2 (from panel E) are depicted. The means of FHL2 overexpressing cells under UF and FHL2-knockdown (UF shFHL2 (+dox)) cells are shown, with the aspect ratio normalised according to clusters 1 and 2. Inset shows a bar graph of the fraction of cells (from UF, DF, UF FHL2 OE, and DF shFHL2) in each predefined cluster, using k-means clustering approach. Wavy arrows depict how expression levels of FHL2 tunes the fraction of focal adherens junctions. N=3 for UF, DF, and N=2 independent experiments for FHL2 OE (n=1179 cells from control UF and DF clustered into cluster 1 or 2, 236 for UF FHL2 OE, 225 for DF shFHL2). **Scale:** (A, F) 20 µm. **Statistical tests**: Unpaired t-test. ns: p value not significant, ****p<0.0001.

We characterised cell junction phenotypes by systematically categorising junction morphologies using VE-Cadherin immunostaining images of *in vitro* endothelial tissues in UF and DF. As described in Fig. 1C as well as previous *in vivo* studies^36^, a major parameter distinguishing UF and DF phenotypes is the cell aspect ratio. Thus, by plotting the cell aspect ratio versus the fraction of linear, reticular or focal adherens junctions, we identify different cell populations that represent the cells in the UF and DF conditions distinctly (Fig. 3B, 3C, 3D). We show that in healthy endothelial tissues under UF, cells exhibited a high proportion of their perimeter marked by linear junctions (∼50%; Fig. 3B), and a very low fraction of discontinuous focal adherens junctions (∼8%; Fig. 3C). On the other hand, cells under DF have very different junction morphologies, with focal adherens junctions dominating the phenotype, occupying ∼60% of the junction perimeter (Fig. 3C). Further, the fraction of the perimeter occupied by reticular junctions was highly similar between cells under UF and DF (Fig. 3D). Such junction phenotypes are consistent with previous studies showing that cells experiencing low shear or static conditions tend to have discontinuous adherens junctions^62,65^. Our results, however, represent a high resolution and quantitative characterisation of the adherens junction morphologies, where the abundance of the different junction classes serves as key readouts of athero-protective UF versus athero-prone DF endothelial tissues. Thus, our findings strongly demonstrate that in endothelial tissues under athero-prone DF, there is a tendency to build focal adherens junctions, which in turn, results in increased tissue permeability (as shown in Fig. 2L).

Next, we sought to classify cells into clusters with the objective of distinctly representing cell populations based on the similarities and differences between each cell/data point. We performed an unsupervised k-means clustering with multiple parameters of cell and junction morphologies measured in endothelial tissues (aspect ratio, linear, reticular, and focal adherens junction fractions). This analysis revealed two strikingly distinct clusters which were nearly identical to grouping by UF and DF conditions (Fig. 3E, S3A). We, therefore, designated these clusters as UF-like Cluster 1 and DF-like Cluster 2 (Fig. 3E).

With this in-depth characterisation of endothelial junction morphologies during flow-sensing, we then explored whether FHL2 upregulation was sufficient to drive athero-prone morphological phenotypes, even when *in vitro* endothelial tissues were subjected to athero-protective UF for 24 h. Following a similar approach as described above, we analysed cell junction morphologies from VE-Cadherin immunostaining images of control and FHL2-overexpressing cells subjected to UF. FHL2-overexpressing cells under UF displayed a significant increase in the formation of discontinuous focal adherens junctions, with a smaller amount of linear cell junctions (Fig. 3F, 3G, 3H). By plotting these junction parameters with respect to the aspect ratio, we identify that the population of cells overexpressing FHL2 are distinct from the control cells. In particular, FHL2 overexpression cells displayed fewer linear junctions (∼16%; Fig. 3F, 3G) and an augmented proportion of focal adherens junctions (∼43%; Fig. 3F, 3H). Moreover, the proportion of reticular junctions between the control and FHL2-overexpressing cells under UF was not significantly different (∼42% and 41% respectively; Fig. S3B). Overall, the effect of FHL2 overexpression in promoting focal adherens junctions is comparable to cellular phenotypes in response to DF.

We then assessed whether the adherens junction morphologies seen in DF could be rescued by FHL2 knockdown. To this end, we generated a doxycycline (dox)-inducible endothelial cell line expressing tet-on shRNA for FHL2 (TeloHAEC shFHL2), where FHL2 protein levels are significantly reduced, as measured by western blotting (Fig. S4A, S4B) and immunofluorescence (Fig. S4C, S4D). Although FHL2 knockdown cells under DF did not alter the cell aspect ratio or alignment (Fig. S4E, S4F, S4G), we observed a significantly higher fraction of the junction perimeter occupied by linear junctions (∼22%; Fig. S4G, S4H) and a much lower fraction of focal adherens junctions (∼29%; Fig. S4G, S4I) compared to control cells under DF. There was also a significant increase in the fraction of reticular junctions in the FHL2-knockdown cells compared to control cells under DF (Fig. S4H, S4J). Altogether, these results suggest that reducing FHL2 expression partially rescues the effect of DF on focal adherens junction morphologies.

Finally, to compare the effects of varying FHL2 expression on the adherens junction morphologies, we plotted the experimental means of all conditions, along with the centroids of Clusters 1 and 2 (Fig. 3I). As indicated by the black arrows, tuning FHL2 levels in the absence of external mechanical input drives phenotypes towards UF-like and DF-like Clusters 1 and 2. Furthermore, with the clustering approach described in Fig. 3E, in each condition, we determined the number of cells in UF-like Cluster 1 and DF-like Cluster 2 (Fig. 3J). Strikingly, in conditions with high FHL2 levels (∼94% of DF, ∼70% of UF FHL2 OE), a majority of cells were found in the DF-like Cluster 2 (Fig. 3J, S3C, S3D), and on the other hand, in conditions with low FHL2, most cells (∼99% of UF) belonged to the UF-like Cluster 1 (Fig. 3J, S3C, S3D). In addition, with FHL2-knockdown under DF, a high fraction (∼59%) of cells reverts to UF-like Cluster 1 (Fig. 3J, S3K, S3L).

Overall, our results demonstrate that FHL2 expression is sufficient to drive athero-prone phenotypes even under UF (Fig. 3K). We also find that reducing FHL2 levels partially reverses athero-prone phenotypes induced by DF flow, by abrogating the amount of leaky focal adherens junctions (Fig. 3K). Taken together, we show that FHL2 expression tunes endothelial function through remodelling of adherens junctions.

### Force-sensitive FHL2 binding to stress fibres is essential for driving athero-prone endothelial phenotypes

Given that FHL2 is enriched on stress fibres in DF or FHL2 overexpression, we investigated whether this localisation arises from its force-dependent recruitment to actin filaments^44,46^. Thus, we abrogated the mechanosensitive localisation of FHL2 to F-actin by generating a doxycycline (dox)-inducible TeloHAEC cell line with four phenylalanine-to-alanine point-mutations^46^, F80A, F141A, F200A, and F263A (TeloHAEC FHL2-F1-4A-mEmerald, F1-1A OE; Fig. 4A, S5A, S5B).

**Figure 4:**
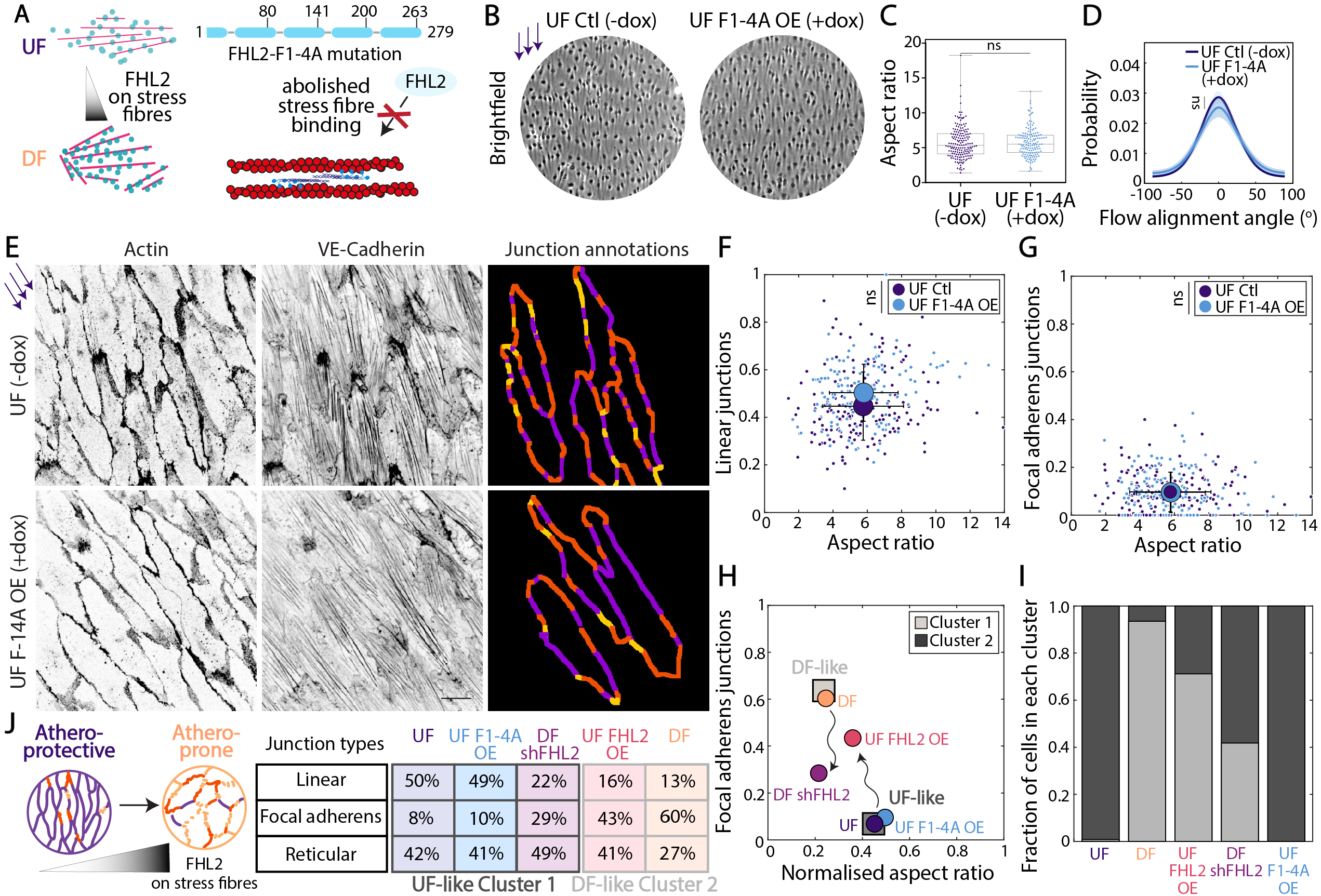
FHL2 binding to stress fibres is essential for promoting athero-prone phenotypes. **(A)** Schematic of the four phenylalanine point mutations created in FHL2, to generate FHL2 F1-4A (mutations previously described in ^46^). **(B)** Inverted contrast brightfield images of TeloHAECs overexpressing mEmerald tagged-FHL2 WT or FHL2 F1-4A. **(C)** Graph shows the aspect ratio (ratio of the major to the minor axis of a cell) of control (-dox) and F1-4A overexpressing (OE; +dox) TeloHAECs under UF and DF. N=2 independent experiments (n=143 for Ctl and 167 for F1-4A OE). **(D)** From brightfield images, the alignment of control and F1-4A overexpressing cells under UF are calculated by local gradient orientation, and represented as the probability distribution of cells with the flow alignment angles (centred around 0°). Shaded error bars representing standard deviation are shown. N=3 independent experiments (n=34 for UF Ctl, and n=39 for UF F1-4A OE; fields of view, with >100 cells/field). **(E)** Inverted contrast confocal images of control and F1-4A overexpressing TeloHAECs under UF, stained for phalloidin and VE-Cadherin. Annotated junctions (three types) are depicted. **(F, G, H)** Graphs showing the aspect ratio (ratio of the major to the minor axis of a cell) vs. the fraction of the cell perimeter occupied by linear junctions (F), focal adherens junctions (G). N=2 independent experiments (n=143 for UF Ctl, 167 for UF F1-4A OE). One data point belonging to UF Ctl (aspect ratio 18.25, linear 0.62, focal adherens 0.01) used in the analyses is excluded from the graph for representation purposes. The UF Ctl means were pooled from 5 experiments for statistical tests. **(I)** Centroids of Clusters 1 and 2 (from Fig. 3) are depicted. The means of F1-4A and FHL2 overexpressing cells under UF, and FHL2 knockdown cells under DF are shown, with the aspect ratio normalised according to Clusters 1 and 2. Inset shows a bar graph of the fraction of cells (from UF, DF, UF FHL2 OE, and DF shFHL2, UF F1-4A OE) in each predefined cluster, using k-means clustering approach. Wavy arrows depict how expression levels of FHL2 tune the fractions of focal adherens junctions. N=3 for UF, DF, and N=2 independent experiments for F1-4A OE (n=1179 cells from control UF and DF clustered into cluster 1 or 2, 236 for UF FHL2 OE, 225 for DF shFHL2, 167 for UF F1-4A OE). In the box-and-whisker plot, the box extends from the 25th to the 75th percentile, the whiskers show the minimum and maximum values, and the line within the box represents the median. **Scale:** (E) 20 µm. **Statistical tests**: (C) Student’s t-test followed by Mann-Whitney test. (D) Kolmogorov-Smirnov test. (F, G) Unpaired Student’s t-test. ns: p value not significant.

In UF conditions, F1-4A-overexpressing cells recapitulated the alignment and aspect ratio observed in control cells (Fig. 4B, 4C, 4D). Further, F1-4A-overexpressing cells had a high proportion of linear (∼49%; Fig. 4E, 4F) and reticular junctions (∼41%; Fig. S5C), and very few focal adherens junctions (∼10%; Fig. 4G), in contrast to the FHL2-overexpressing cells under UF (Fig. 2I, 2J, 2K). Overall, there were no significant differences in the junction morphologies upon F1-4A overexpression, and such junction types strongly resembled control cells under UF (Fig. 4E, 4F, 4G). This suggests that mutating the residues important for force-activated binding of FHL2 to actin abrogates its role in promoting an athero-prone phenotypes.

Finally, to compare the effects of stress fibre-bound FHL2 and varying FHL2 levels on junction morphologies, we depicted the mean of F1-4A-overexpressing cells on the graph shown in Fig. 3I. Here, F1-4A-overexpressing cells were strikingly similar to UF conditions (Fig. 4H). We further determined the fraction of F1-4A-overexpressing cells in each cluster and found that all the cells belonged to the UF-like cluster 1 (Fig. 4I). These results indicate that the overexpression of cytoplasmic, non-stress fibre binding mutant FHL2 had no detrimental effects on cell aspect ratio, alignment, or junction morphologies when subjected to UF (Fig. 4J). This also implies that in cells that do not overexpress actin-binding FHL2, the cellular phenotypes mimic control cells subjected to UF. Overall, taking together our previous findings, we demonstrate a crucial function for FHL2-binding to stress fibres, by showing that this force-sensitivity is necessary for promoting an athero-prone phenotype.

### FHL2 promotes actomyosin contractility

Adherens junctions are coupled to the actomyosin cytoskeleton; as such, they are strongly modulated by changes in RhoGTPase signalling^29,61,66–68^. For instance, in endothelial cells, focal adherens junctions arise at the ends of actomyosin stress fibres^29,63^. Thus, we sought to explore whether changes in FHL2 expression modulates cellular contractility.

First, we measured phosphorylated myosin light chain (pMLC) in control and FHL2-knockdown cells. Immunostaining for pMLC and actin revealed a ∼30-40% reduction in FHL2-knockdown cells (Fig. 5A, 5B). These measurements were corroborated with western blotting, where a 30% reduction in pMLC was observed in FHL2-knockdown cells (Fig. 5C, 5D, S6A, S6B). Next, by employing traction force microscopy (TFM), we computed cellular force generation and the overall strain energy exerted by cells onto a compliant fibronectin-coated substratum. We observed that FHL2-knockdown resulted in 2-fold lower energies compared to control cells (Fig. 5E, 5F). These findings underscore the importance of FHL2 in promoting Rho-mediated actomyosin contractility. This also suggests that FHL2-mediated contractility is essential for discontinuous adherens junction morphologies, characteristic of endothelial dysfunction in athero-prone tissues (Fig. 5G).

**Figure 5:**
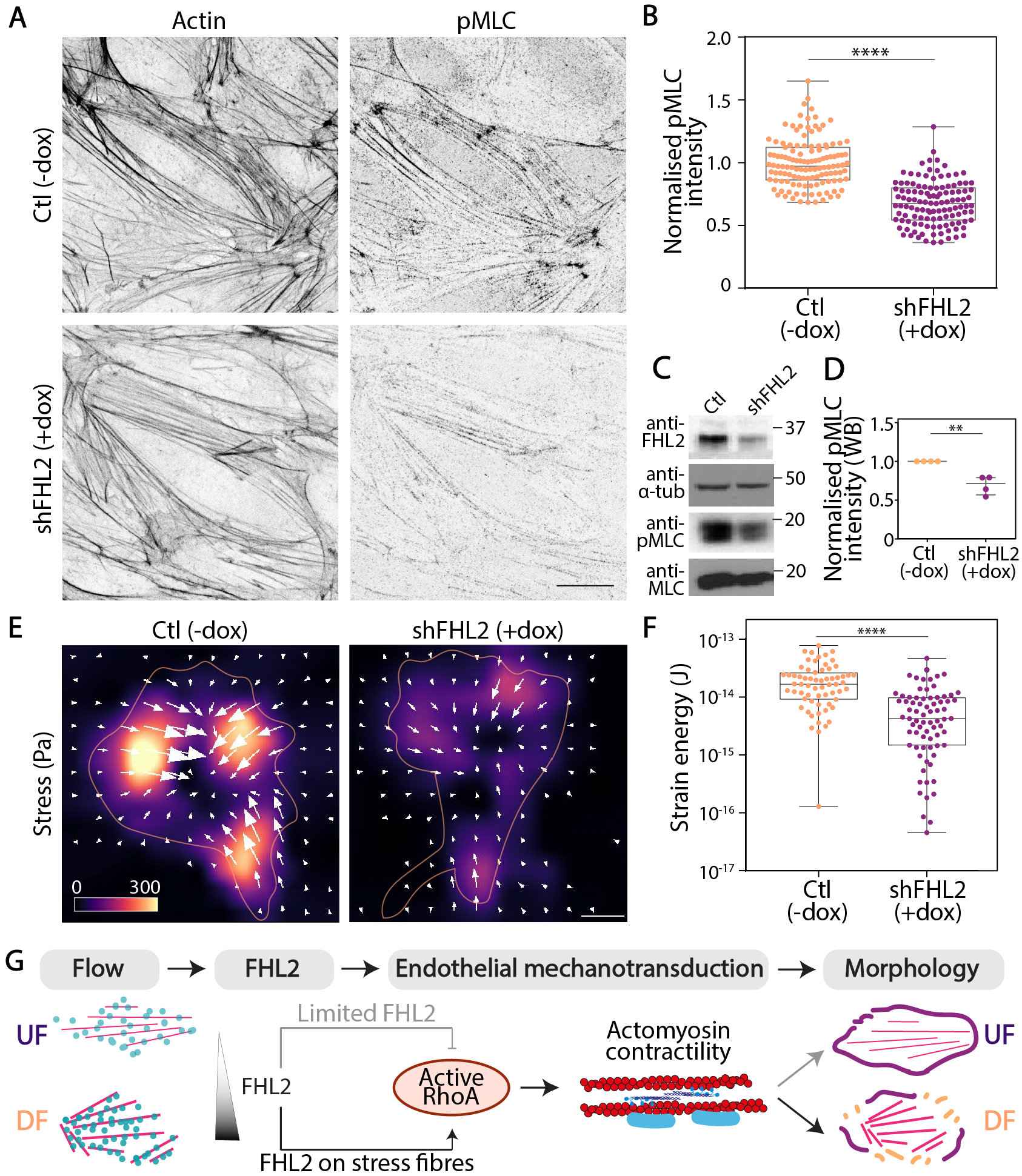
FHL2 expression promotes actomyosin contractility. **(A)** Inverted contrast images of TeloHAECs immunostained for phalloidin (to label actomyosin stress fibres) and phosphorylated (serine 19) myosin light chain (pMLC). **(B)** Graph shows the normalised pMLC levels (pMLC intensity/actin intensity) in control (Ctl; -dox) and FHL2-knockdown cells (shFHL2; +dox). N=3 independent experiments (n=125 for Ctl, 119 for shFHL2). **(C)** Western blot of lysates from Ctl and shFHL2 cells. Samples were analysed by immunoblotting for FHL2, pMLC, MLC, and tubulin (loading control). **(D)** Normalised pMLC intensity (pMLC/MLC intensity) in Ctl and shFHL2 cells. N=4 independent experiments. Dots represent experimental means, whiskers show the minimum and maximum values, and the line within represents the median. **(E, F)** Traction stress maps (D) and strain energy of each cell (in Joules; (E)) of Ctl (-dox) and FHL2-knockdown (shFHL2; +dox) cells shown. Vectors indicate direction and magnitude of the stress. N=2 independent experiments (n=64 for Ctl, 75 for shFHL2). **(G)** Schematic showing that the levels of FHL2 in cells alters actomyosin contractility. Taken together with Fig. 3 and 4, schematic shows that the localisation of FHL2 to stress fibres is important for endothelial mechanoresponses. **Scale:** (A) 50 µm, (E) 10 µm. **Statistical tests**: (B, E) Student’s t-test followed by Mann-Whitney Test. **p<0.01, ****p<0.0001.

### FHL2 promotes actomyosin contractility through crosstalk with the microtubule cytoskeleton

RhoA activity is regulated by Rho guanine exchange factors (RhoGEFs) or RhoGTPase activating proteins (RhoGAPs)^69^. The RhoGEF, GEF-H1, is one of the most abundantly expressed GEFs in endothelial cells and has been implicated in cell mechanotransduction^70–74^. When GEF-H1 is sequestered on microtubules, it is inactive. When GEF-H1 is released into the cytoplasm, it renders RhoA active^70,75^.

We first tested whether FHL2 alters GEF-H1 localisation and activity. In control cells, GEF-H1 localisation was primarily cytoplasmic (Fig. 6A), suggesting active GEF-H1. We observed high intensity peaks of tubulin, with the lack of GEF-H1 corresponding to these peaks in control cells (Fig. 6B). Similarly, we were able to quantitatively assess the amount of GEF-H1 on a microtubule versus in the cytoplasm (Fig. 6C). By contrast, FHL2 knockdown resulted in enhanced microtubule-bound GEF-H1 (Fig. 6A, 6B). As shown in the intensity profiles, GEF-H1 peaks are coincident with tubulin, indicating a strong localisation of GEF-H1 to microtubules in FHL2-knockdown cells. FHL2-knockdown also increased the microtubule-bound GEF-H1 (Fig. 6C). Furthermore, we quantified the relative amount of GEF-H1 bound to microtubules versus free in the cytoplasm to confirm that the ratio of microtubule- to cytoplasmic GEF-H1 was higher in FHL2 knockdown cells (Fig. 6C).

**Figure 6:**
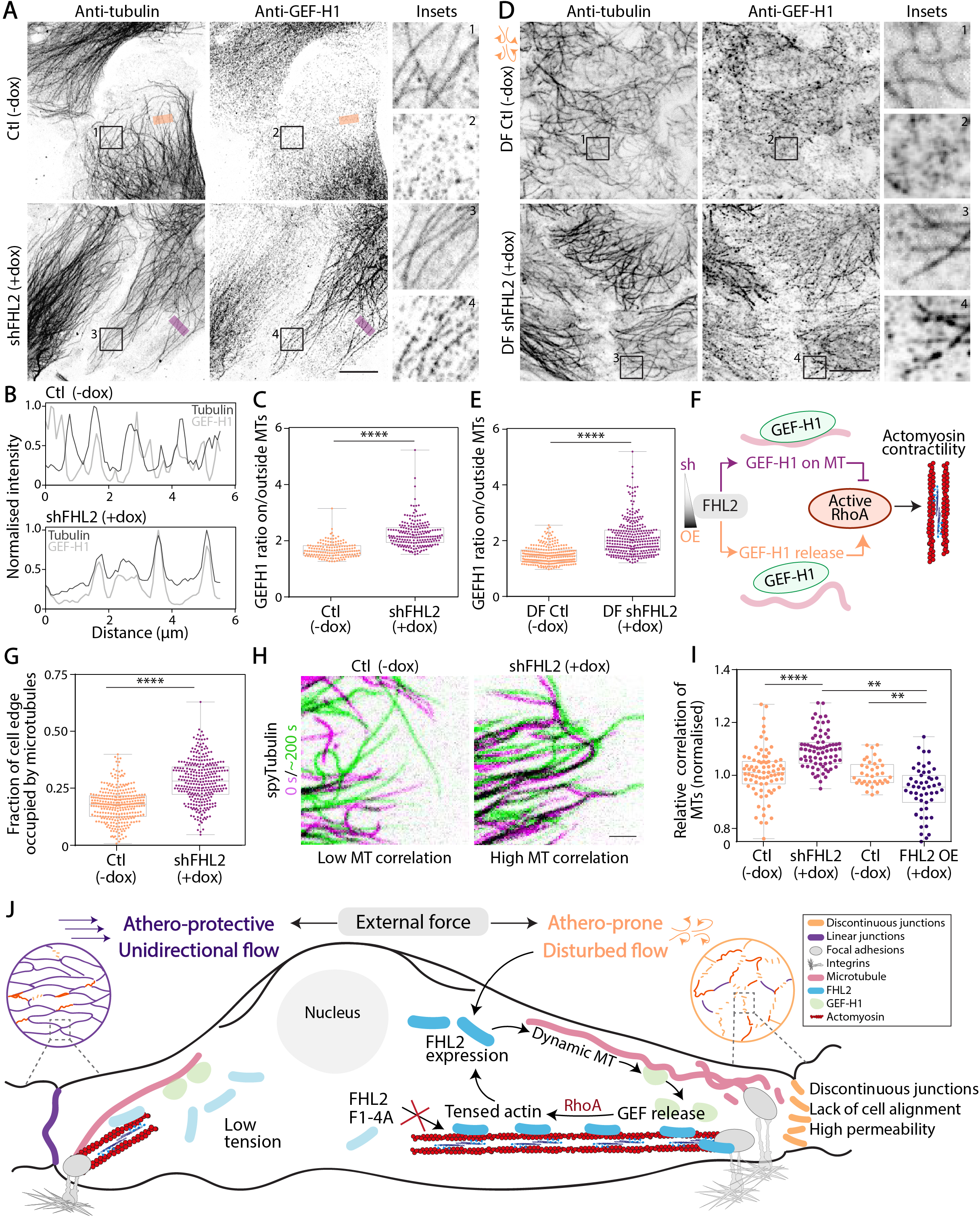
FHL2 promotes the release of GEFH1 by augmenting microtubule dynamics. GEF-H1 sequestered on microtubules renders the GEF inactive, while cytoplasmic localisation (released from microtubules) marks the active pool of GEF-H1 that can activate RhoA. **(A)** Inverted contrast confocal images of control (Ctl; -dox) and FHL2-knockdown (shFHL2; +dox) TeloHAECs, immunostained for α-tubulin and Rho guanine nucleotide exchange factor, GEF-H1. **(B)** Line scans showing GEF-H1 and tubulin intensities from a representative region of a microtubule. **(C)** Graph showing the ratio of GEF-H1 intensity on vs. off microtubules. N=2 independent experiments (n=151 for Ctl, 203 for shFHL2; each dot represents a region of interest; ∼2 regions per cell measured). **(D)** Inverted contrast confocal images of Ctl (-dox) and shFHL2 (+dox) TeloHAECs under DF, immunostained for α-tubulin and Rho guanine nucleotide exchange factor, GEF-H1. **(E)** Graph showing the ratio of GEF-H1 intensity on vs. off microtubules. N=2 independent experiments (n=262 for DF Ctl, 305 for DF shFHL2; each dot represents a region of interest; ∼2 regions per cell measured). **(F)** Schematic showing that FHL2 expression promotes the release of GEF-H1 into the cytoplasm, which can activate RhoA. On the other hand, cells with low FHL2 levels exhibit microtubule-bound inactive GEF-H1. **(G)** Graph shows the fraction of the cell edge occupied by microtubules, by measuring α-tubulin intensity in regions marking the cell protrusions. N=3 independent experiments (n=312 for Ctl, 307 for shFHL2; each dot represents a regions; ∼1-2 regions per cell measured). **(H)** Using spyTubulin labelling in cells imaged over time, microtubule fluctuations between each frame are represented as the correlation index. Inverted contrast images of microtubules, where two representative frames, approximately 200s apart are shown in green and magenta. A high correlation index corresponds to low fluctuations or dynamics, while a low correlation index suggests a highly fluctuating microtubule network. **(I)** Graph represents the normalised relative correlation of microtubules over time in Ctl, shFHL2 (+dox), and FHL2 OE (+dox) cells. N=3 independent experiments for Ctl and shFHL2, 5 independent experiments for Ctl and FHL2 OE (n=79 for Ctl, 74 for shFHL2, 37 for Ctl, 57 for FHL2 OE; each dot represents a region in a cell protrusion). **(H)** Model of FHL2-mediated endothelial mechanoresponses. Upregulation of FHL2 and its binding to stress fibres in response to DF results in the release of GEF-H1 from microtubules into the cytosol, where it can activate RhoA. In turn, cells exhibit hypercontractility, where stress fibres build tension and pull adherens junctions to cause breakages and discontinuity. Subsequently, with discontinuous focal adherens junction formation, endothelial tissues with high FHL2 have increased permeability and tissue dysfunction. **Scale:** (A, D) 10 µm, (H) 2 µm. **Statistical tests**: (C, E, G) Student’s t-test followed by Mann-Whitney test, (H) One-way ANOVA followed by Tukey’s multiple comparisons test. ****p<0.0001.

Next, we explored whether FHL2-dependent GEF-H1 release is responsible for athero-prone phenotypes. In control cells under DF, GEF-H1 is predominantly cytoplasmic, as shown by the lower microtubule-bound GEF-H1 (Fig. 6D, 6E). Strikingly, we observed that GEF-H1 is sequestered on microtubules in FHL2-knockdown cells even under DF (Fig. 6D, 6E). All these findings provide a mechanism by which FHL2 promotes contractility in endothelial tissues, through the release of GEF-H1 from microtubules to activate RhoA (Fig. 6F).

We then investigated the mechanisms by which FHL2 promoted the release of GEF-H1 from microtubules. GEF-H1 release primarily occurs at the cell protrusions and edges through microtubule dynamics, depolymerisation or uncoupling from adhesions^72,76–80^. First, we tested whether FHL2 alters GEF-H1 release through changes in the microtubule organisation. In this case, we selected regions in the cell periphery and quantified the fraction of the region occupied by microtubules. By immunostaining for α-tubulin, in control cells with high FHL2 levels, we observed that significantly fewer microtubules reached the cell edges, as represented by the low fraction of cell edge or protrusion occupied by microtubules (Fig. 6G). On the other hand, in FHL2 knockdown cells, microtubules extended to the cell periphery, with a significantly higher fraction of the cell protrusion occupied by the microtubule network (Fig. 6G). This strongly suggests that FHL2 restricts the microtubule network from reaching the periphery by possibly promoting microtubule depolymerisation close to the edge.

To further provide explanation for FHL2-mediated GEF-H1 release, we analysed fluctuations and dynamics of the microtubule network. By live imaging microtubules (labelling tubulin using Spy-tubulin) and FHL2 (using FHL2-OE mEmerald), we investigated whether FHL2 expression levels altered microtubules dynamics (Fig 6H; movie 2). FHL2-decorated actomyosin stress fibres restrict the penetration of microtubules through them, thus dynamically regulating the fraction of microtubules reaching the cell periphery (movie 4). This is in line with our findings from Fig. 6G that fewer microtubules ventured to the periphery in cells with higher FHL2 levels (control, FHL2 OE). We then tested whether tuning FHL2 levels influenced microtubule dynamics. To this end, we measured the relative correlation between microtubules over time and we found that the microtubule network, upon FHL2 knockdown, displayed fewer fluctuations or variations compared to control cells. This is represented by the higher correlation seen between microtubules across multiple frames of the movie (Fig. 6H, 6I; movie 2). In contrast, with higher FHL2 levels in the overexpression conditions, we also observed a lower microtubule correlation over time, suggesting that increased FHL2 expression renders more dynamic microtubules (Fig. 6I; movie 3). Our results demonstrate that tuning FHL2 expression levels results in altered GEF-H1 activity, where high FHL2 promotes the release of GEF-H1 through its effects on microtubule dynamics. Furthermore, taken together with findings from Fig. 4 and 5, we highlight the crucial role for stress-fibre bound FHL2 in promoting contractility through RhoGTPase signalling to drive athero-prone phenotypes.

## Discussion

We uncover a positive mechanochemical feedback mechanism between transcription and subcellular FHL2 localisation to tune actomyosin contractility via GEF-H1. This positive feedback loop drives hypercontractility of tissues during endothelial flow sensing, along with morphological and functional effects on discontinuous adherens junctions, lack of cell alignment, and high tissue permeability; all of these are classic hallmarks of endothelial dysfunction (Fig. 6J). Here, we show how force-sensitivity of FHL2 regulate endothelial mechanotransduction from the subcellular to tissue-scale morphology and function, in response to one mechanical cue seen in arteries, i.e., flow. Further work is needed to explore how this could be a general phenomenon in response to other external forces (e.g., matrix stiffness, stretch or curvature).

Our results demonstrate that the transcriptional regulation of FHL2 occurs as a very early mechanoresponse to changes in flow and precedes the atherosclerotic disease phenotype. Several studies have focused on the transcriptional changes in healthy vs. atherosclerosis arteries, where transcription factors including KLF2, NF-kB and c-Jun have been largely implicated in disease progression. In addition to known transcription factors and cardiovascular genes previously discovered, we find that over 31 LIM domain proteins, including FHL2, are significantly altered during flow-sensing in endothelial tissues. It is noteworthy that major endothelial transcription factors like KLF2 and KLF4, similar to LIM domain proteins, also contain Zinc finger motifs and bind DNA and are crucial for endothelial tissue function^81,82^. Taken together, this suggests that the LIM domain protein family might be crucial for differentially contributing to mechanotransduction in various other cardiovascular diseases that are primarily mediated by changes in flow, either through their transcriptional regulation or interactions through Zn-finger motifs. Moreover, it is possible that FHL2 primes specific regions in the artery to endothelial dysfunction and subsequent atherosclerotic plaque formation. In this context, it would be also interesting to explore the extent to which perturbed tissue phenotypes are reversible by tuning the transcription of mechanosensitive genes, in this case, FHL2 through its known regulators MRTF/SRF^83,84^. Furthermore, the heterogeneity in FHL2 expression across various tissue types suggests that the protein is differentially regulated across tissues and is modulated to maintain specific functionality. Thus, the distinct expression levels as well as localisation indicate that FHL2 could be a useful biomarker of the state of endothelial tissues, particularly as it is a crucial regulator of endothelial mechanoresponses.

We identify a function of the stress fibre-binding ability of FHL2. Although the mechanosensitive binding of FHL2 to stress fibres has been well described^44,46,48^, the function of this binding was unknown. We show that the force-sensitive binding of FHL2 to actomyosin stress fibres promotes RhoA activation via GEF-H1 that, in turn, enhances contractility. We speculate that the concentration of cytoplasmic and stress fibre-bound FHL2 (possibly through nuclear-to-cytoplasmic shuttling or transcriptional status^48,83^) can serve as a way to tune this positive feedback. Thus, this provides a pathway by which overall FHL2 transcriptional control coordinated with local subcellular mechanotransduction regulate RhoGTPase signalling.

Our results strongly suggest that FHL2 mediates force-dependent actomyosin-microtubule crosstalk. We show that FHL2-enriched stress fibres promote microtubule dynamics and prevent the microtubule network from reaching the cortex. In this case, local microtubule depolymerisation, instability at the cortex, or uncoupling from integrin-mediated focal adhesions can cause GEF-H1 release into the cytoplasm^70,72,77,80^. It is possible that FHL2, in addition to its force-sensitive binding to actin, interacts with focal adhesion components or microtubule-actin crosslinking factors to tune GEF-H1 localisation and activity^85–88^. Furthermore, in the context of endothelial tissue function, our findings are in agreement with previous work showing that activated GEF-H1 and microtubule depolymerisation drive vascular permeability in both *in vitro* and *in vivo* models^73,89–92^.

Endothelial function is a multi-scale process that requires coordination of temporal events that control and maintain health. For instance, tissue-scale permeability is a function of cellular mechanotransduction which occurs at the junctions, and different adherens junction morphologies contribute to this mechanoresponse. In various tissue types, adherens junctions are mechanosensitive and are coupled tightly to the actomyosin network through force-dependent recruitment of proteins such as vinculin or α-catenin^18,19,22,25,29,66,93–97^. In endothelial tissues, discontinuous focal adherens junctions are described as points of high tension^61–63^. Our work suggests a mechanism where FHL2 binding to stress fibres increases force generation to drive adherens junction remodelling. It is tempting to speculate that, through its effects on contractility, FHL2 directly or indirectly alters the activity of a key mechanosensitive protein at adherens junctions, resulting in endothelial physiology or dysfunction as described in this study. We speculate that the physiological role of FHL2 in mediating force-adaptation and endothelial permeability might be important for processes like immune response or inflammation. Finally, it would be interesting to explore how the transcriptional regulation of FHL2 mediates adherens junction remodelling, and whether this can be tuned to engineer endothelial physiology. Overall, our work bridges subcellular mechanochemical feedback and transcriptional regulation to tissue-scale endothelial mechanoresponses.

## Acknowledgements

We thank Jiayu Zhu (University of Chicago) for expert advice on flow setup, Benoit Vianay (CEA, Grenoble) for traction force analysis methods, and Luisa Arispe (Northwestern University) and Kashmeera Baboolall (University of Chicago) for discussions. We also thank the Genomics Facility, DNA Sequencing Facility and the Flow Cytometry Core at the University of Chicago. S.S. received support from the American Heart Association (#915248), and Schmidt Sciences, LLC. M.L.G. acknowledges support from NIH RO1 R01GM143792.

## Author contributions

Conceptualization: S.S. and M.L.G.; experimental design and methodology: S.S., J.D., H.R.K., S.T-P.; analysis: S.S., J.D., E.B., T.C., W.H.C.; visualization: S.S., E.B.; investigation: S.S.; resources: S.S., J.D.; interpretation: S.S., M.L.G; writing: S.S. and M.L.G, with input from all authors; supervision: M.L.G and Y.F.; funding acquisition: S.S. and M.L.G.

## Data Availability

Plasmids will be deposited to Addgene prior to publication. RNA sequencing data will be deposited at the Gene Expression Omnibus prior to publication. Analysis codes will be published on an open-source platform or provided upon request.

## Materials and Methods

### Cell culture, treatments and transfection

Human aortic endothelial cells (teloHAECs) were provided by Yun Fang (University of Chicago), who purchased them from ATCC (#CRL-4052). Cells were grown at 37°C with 5%CO_2_ using EGM-2 medium supplemented with SingleQuots from Lonza (CC-3156 & CC-4176). Cells were passaged using 0.25% trypsin EDTA every 2–3 days. Cells were tested for mycoplasma every 6 months or as necessary (no mycoplasma detected) using InvivoGen MycoStrip^TM^ (rep-mys-10) and spyDNA/Hoescht staining. Tet-inducible gene expression was carried out with 200ng/ml doxycycline in all indicated experiments (+dox). Cells were electroporated using the Neon™ Transfection System, with 1 pulse at 1350V for 30ms, and plated on appropriate supports, coverslips or culture dishes as needed. The growth medium was replaced 8 or 24 h post-transfection. For treatment with Rho-kinase (ROCK) inhibitor Y-27632, cells were treated with 10 µM for 4 h. To visualize microtubules and actin during live-cell imaging, cells were incubated with Spytubulin (CY-SC203; 1:2000) and Spyfastact (NC2204057; 1:2000) for 1-2 h, and fresh growth medium was replaced prior to imaging. For traction force experiments, cells were incubated with CellMask Orange (Invitrogen, Cat# C10045; 1:5000) for 20 min, washed once and replaced with fresh growth medium prior to imaging.

### Cell line generation

FHL2 overexpression and knockdown teloHAEC cell lines were made by CRISPR-mediated knock-in to the AAVS1 locus of tet-inducible FHL2-mEmerald or a tet inducible mir30a shRNA containing two previously published sequences targeting FHL2 (see below). These sequences were ordered from IDT and cloned into a previously published vector (pMK243) with homology arms targeting the human AAVS1 locus and co-expression of tet-on3g. mir30a based shRNA was designed as previously described^98^. Each plasmid was co-transfected with a plasmid encoding CAS9 and a sgRNA targeting the AAVS1 locus (AAVS1 T2 CRIPR in pX330) at a ratio of 1:1 using the neon transfection system. All constructs were sequenced at the University of Chicago Genomics core and verified prior to use. One week after transfection, cells were induced with doxycycline and sorted for expression of GFP or CFP. Cells were induced and sorted as needed before use for experiments. For flow experiments, cells were re-sorted to ensure a good population of positive GFP or CFP cells.

#### FHL2 shRNA sequences

FHL2 shRNA 1: CGAGACTTTCTTCTAGTGCTTT ^99^

FHL2 shRNA 2: AAACGAATCTCTCTTTGGCAAG ^100^

#### Primers used in this study

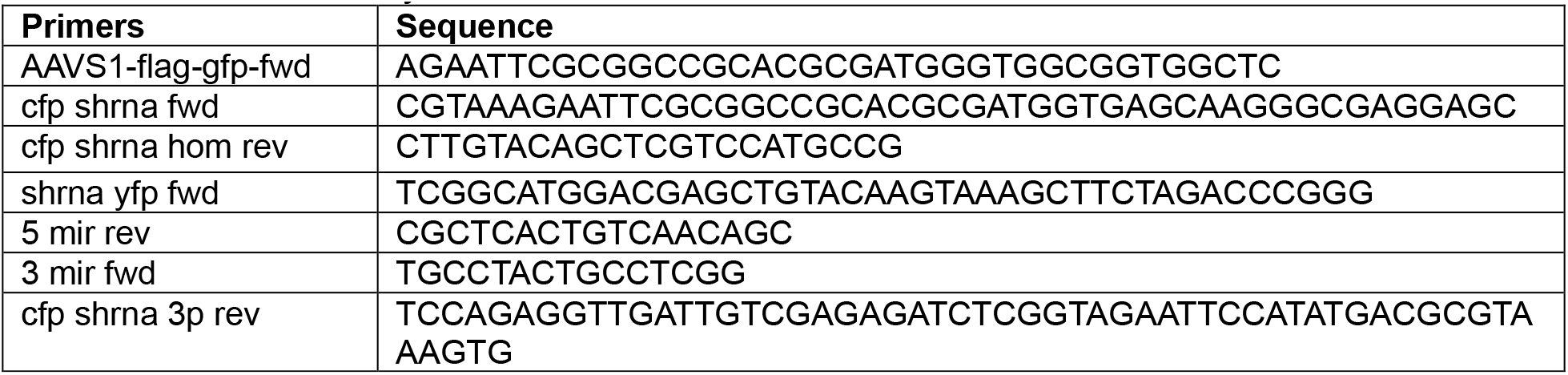

### Flow experiments

Cells were cultured in 6-well plates with EGM-2 medium supplemented with 4% dextran (Sigma-Aldrich, St. Louis, MO, 31392). Hemodynamic flows were applied to using a 1o tapered stainless steel motorised cone (UMD-17 (Arcus20 Technology, Livermore CA). The rotation of the cone captures healthy unidirectional flow (UF) seen in athero-protective distal internal carotid artery, while the disturbed flow (DF) mimics athero-susceptible human carotid artery^33^. During flow application to cells, the flow device was placed at 37°C with 5%CO_2_.

### RNAsequencing

Cells subjected to UF and DF were lysed, and total RNA was collected using a NucleoSpin RNA kit (Macherey-Nagel, #740955). Two technical replicates for each condition were pooled together to obtain an experimental replicate. The RNA samples from three separate experimental replicates were submitted to the University of Chicago Genomics Facility. The quality and quantity of RNA were assessed using the Agilent bio-analyzer (RNA Integrity Number from 9.9 to 10). Using a TruSEQ mRNA RNA-SEQ library protocol (Illumina provided), strand-specific RNA-SEQ libraries were prepared. Libraries were then sequenced using Illumina NovaSEQ6000 (illumina provided reagents and protocols) with ∼60M PE reads/sample. The reads were aligned with the Homo Sapiens genome hg19/GCh37 reference by psudeoalignment using Kallisto 0.46.1. All raw files and analysis will be uploaded on Gene Expression Omnibus prior to publication.

### Differential gene expression analysis

DESeq2 (version 1.40.2) in R software (version 4.3.1) was used to normalize read counts per transcript and perform differential gene expression analysis. DESeq2 normalizes the data according to the following steps: 1. calculation of a fictive reference sample defined as the geometric mean for each gene across all samples, 2. counts for each gene for each sample are divided by this reference value, 3. size factors are estimated by calculation of the median of these ratios across samples, 4. raw gene counts for sample are divided by the sample size factor to obtain normalized counts. This method normalization accounts for library size differences and samples bias. DESeq2 uses normalized counts to calculate log2fold changes and p-values using the Wald test. The Benjamini-Hochberg correction was performed to obtain p-values adjusted for multiple testing. Genes were defined as significantly differentially expressed at an adjusted p-value of < 0.05.

### Gene set enrichment analysis

Simple gene set enrichment analysis was performed in R software using clusterProfiler 4.8.3 ^101^ and version 7.5.1 of the molecular signature database^102^. Differentially expressed genes were tested for overrepresentation in Gene Ontology resource (release date 2023-01-01) clusters (^103^ and the Gene Ontology Knowledge Base, 2023) of molecular function, cellular compartment and biological processes. The HGNC gene nomenclature committee annotation of genes (https://www.genenames.org/) was used as an additional database to test in what gene families selected genes were overrepresented using clusterProfiler.

### Partial carotid ligation surgery

All mouse procedures were approved by the Institutional Animal Care and Use Committee (IACUC) of The University of Chicago. Partial carotid artery ligation was surgically performed to introduce acute disturbed flow activating endothelial cells in the left carotid artery. Briefly, male 7-9 weeks old C57BL/6 mice (The Jackson laboratory) were anesthetized by intraperitoneal injection of ketamine (100 mg/kg) and xylazine (10 mg/kg) mixed solution. A ventral midline incision (∼ 5 mm) in the neck was opened by a tiny scissor (F.S.T 14090-11). The left carotid artery was exposed by blunt dissection under microscopy. Then, the left external carotid, internal carotid, and occipital artery were ligated with 6-0 silk suture while the superior thyroid artery remains intact. After that, the incision was closed with Nylon monofilament suture (Air-Tite products) and the cut was smeared with Betadine to prevent infection. 48 hours after ligation, the animals were euthanized, and both carotid arteries were harvested after saline perfusion via the left ventricle after severing the inferior vena cava. After euthanasia and saline perfusion, carotid arteries were isolated en bloc. Carotid arteries were fixed in 4% paraformaldehyde, sequentially immersed in 30% sucrose and optimal cutting temperature compound (OCT):30% sucrose (1:1) mixed solution, and then embedded in OCT. Frozen-embedded samples were sectioned in 5- to 8-μm thickness. Sections were then immunostained as described later in the Methods Section.

### Intimal RNA isolation from carotid arteries

After careful isolation, the carotid lumen was quickly flushed with 350 μl of QIAzol lysis reagent (QIAGEN, MD, USA) using a 29-gauge insulin syringe, and the elute was collected in a microfuge tube. The elute was further applied for intimal RNA isolation. Total RNA was reverse transcribed with Maxima First Strand cDNA Synthesis Kit (Thermo Fisher Scientific, MA, USA) and SYBR Green qRT-PCR was performed.

### Quantitative real-time PCR (qPCR)

Total RNA was extracted with GenElute Mammalian Total RNA Miniprep Kit (Sigma). 0.2 μg mRNA was reverse-transcribed using High-Capacity cDNA Reverse Transcription Kit (ThermoFisher). The cDNA was amplified by LightCycler 480 II (Roche) system using a SYBR Green probe. Absolute quantification of the gene of interest was normalized to the geometric mean of β-actin, GAPDH and Ubiquitin.

#### Primers used in this study

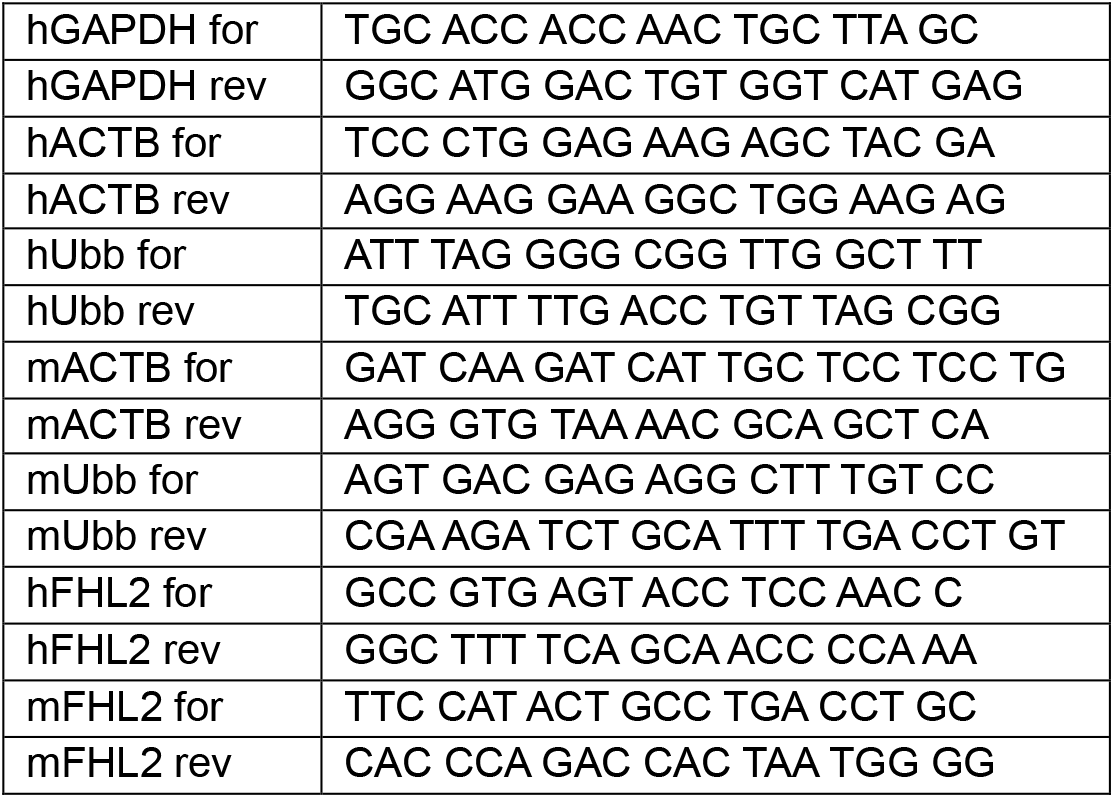

### Immunostaining

Endothelial cells were washed with 1x PBS gently, and subsequently fixed using one of the following 3 methods: (1) warm 4% PFA for 10 min at 37°C, followed by 0.2% Triton X-100, (2) ice-cold methanol for 2-3 min at -20°C to stain for microtubules and GEF-H1, (3) 4% warm PFA + 0.2% glutaraldehyde + 0.25% Triton X-100 for 10 min at 37°C. In the case of method 3 using PFA and glutaraldehyde, the free aldehyde groups are quenched with a 1 mg/ml sodium borohydride freshly added to cytoskeletal buffer (10 mM MES or MOPS, 150 mM NaCl, 5 mM ethylene glycol-bis(β-aminoethyl ether)-N,N,N′,N′-tetraacetic acid (EGTA), 5 mM MgCl_2_ and 5 mM glucose, pH 6.1) and placed on ice or at 4°C for 10 min. After fixation, a 5% bovine serum albumin solution in PBS was used for blocking at room temperature for 1 h. The same BSA solution was used to prepare the primary and secondary antibody solutions, which were incubated for 1 h each, with PBS washes in between the incubation periods. In the case of flow experiments, primary antibody incubation was carried out at 4°C overnight. Coverslips were mounted on glass slides using Prolong Gold Antifade Mounting medium.

#### Antibodies used in this study

FHL2 (rabbit, Sigma, HPA006028, 1:100), VE-Cadherin (mouse, Santa Cruz, SC-9989, 1:2000) pMLC (rabbit, Cell Signaling, 3671S, 1:1000), GEF-H1 (rabbit, Abcam, ab155785, 1:500), alpha-tubulin (Y1/2, rat, Abcam, ab6160). Secondary antibodies were purchased from Jackson Antibodies.

### Western blotting

Cells were lysed using Laemmli buffer (60 mM Tris-HCl pH 6.8, 10% glycerol, 2% sodium dodecyl sulfate (SDS) and 50 mM dithiothreitol (DTT)), with the addition of anti-protease and phosphatase inhibitors. Cell lysates samples were boiled for 5 min at 95 °C before freezing at -20°C or loading on to PAA gels. Proteins were transferred onto a nitrocellulose membrane, at 100 V for 1 h. Membranes were blocked with 5% milk in Tris-buffered saline, 0.1% Tween 20 detergent (TBST) for 1 h, followed by primary antibody (diluted in a solution of 5% milk in TBST) overnight at 4°C. Membranes were then washed with TBST and incubated with horseradish peroxidase (HRP)-conjugated secondary antibody (diluted in 5% milk in TBST) for 1 h at room temperature. Bands were revealed using ECL chemoluminescent substrate (Thermo Fischer Pierce ECL western blot substrate).

#### Antibodies used in this study

GAPDH (rabbit, G9545, 1:2000), other antibodies were the same as for immunostaining and are listed above. Secondary antibodies used were anti-rat-HRP (Thermo Fischer, A10549, 1:10000), anti-rabbit HRP (Cell Signaling, 7074S, 1:10000), anti-mouse HRP (Cell Signaling, 7076, 1:10000).

### Permeability assay

Cells were plated to confluency in 0.4 µm 12-well transwell plates (Corning, #734-1579). Total volume in the top chamber was 200 µl, and 1.5 ml of media was added to the bottom chamber. Cells were cultured on transwell inserts for 24-48 h until they formed a confluent monolayer. The media from the top chamber was carefully aspirated and replaced with 200 µl media containing 10 µl/ml FITC-dextran (Sigma, FD4-100MG). Media from the bottom chamber was replaced with fresh media. Cells in transwells were incubated for 8 hours in 37°C with 5%CO_2_. After 8 h, media from the top and bottom chambers were collected in a centrifuge tube for processing. 50 µl each from the top and bottom chambers were transferred into separate wells in a 96 well plate. A standard curve with FITC-dextran was created and the fluorescence was measured with an excitation wavelength of 488 nm and emission of 520 nm, using a plate reader. The data was analysed according to previous literature^104^. Briefly, using the volume of media in each chamber, time, surface area/well, and the measured fluorescence intensity in each condition, we calculated the fold change in permeability after calculation of the absolute concentration of FITC in the top and bottom chambers.

### Traction force experiments

Coverslips were silanised for 10 min in a solution of 1% (v/v) 3-(trimethoxysilyl)propyl methacrylate and 1% (v/v) acetic acid in ethanol, and washed twice with absolute ethanol and dried on Whatmann paper. PAA gels of 8.4 kPa Young’s Modulus were prepared using 3.125 ml 40% acrylamide (Bio-Rad Laboratories, Hercules, CA), 0.833 ml 2% bis-acrylamide (Bio-Rad), and 1.042 ml water. To a 500 µl aliquot of this gel mixture, 5 µl of 110-nm sulfate-modified fluorescent microspheres (Invitrogen, Carlsbad, CA). Following this, 2.5 µl of 10% ammonium persulfate (APS) and 0.25 µl of tetramethylethylenediamine (TEMED) were added and the gel solution was mixed well. A 14 µl drop of the solution was placed on each silanised coverslip (22 × 22 mm) and immediately a slide or 18 × 18 mm coverslip was placed gently over the solution. The solution was allowed to polymerize for 30 min at room temperature. Once the gel polymerised, Milli-Q water was added over the coverslips, and the top glass was detached using a scalpel blade. The polymerized gel was then activated for 5 min under ultraviolet light with Sulpho-SANPAH and washed with 10 mM HEPES (4-(2-hydroxyethyl)-1-piperazineethanesulfonic acid) twice. Gels were coated with a fibronectin solution in PBS immediately after washing and placed in the 37°C incubator. Excess matrix protein was washed with PBS and 25000 cells were plated per coverslip and allowed to adhere overnight. Right before the traction force experiments, cells were stained with CellMask Orange to visualise the cells and membranes and rinsed two times with warm PBS. Samples were then mounted on a Chamlide Magnetic Chamber and placed in a stage-top incubator maintained at 37°C and 5% CO_2_. Z-stacks of the fluorescent beads and cell mask were imaged.

### Fluorescence microscopy

Flow and traction force experiments were imaged on a Nikon spinning disc confocal microscope (Nikon TI-E, Nikon, Tokyo, Japan). Images were acquired on Andor Zyla 4.2 CMOS camera (Andor Technology, Belfast, UK), using 642, 561, and 488 lasers. All other imaging was done using a point scanning confocal microscope (Zeiss Airyscan LS980), equipped with laser lines at 405, 491, 561, 642.

### Imaging and image analysis

All imaging settings were maintained constant between conditions that are compared directly. ImageJ and MATLAB were used for image analysis. All codes will be made publicly available or provided upon request from authors.

#### Cell alignment

Brightfield images of *in vitro* endothelial tissues under flow were acquired using a tissue culture microscope. Each frame containing several hundred cells were divided into 2 or 4 images for measuring orientation in small fields of view. Using the ‘local gradient orientation’ in the directionality function in Fiji. Briefly, a local gradient orientation derived by using a 5x5 Sobel filter is then used for the histogram. The flow alignment angle (o) is the direction angle which represents the centre of the Gaussian. This has been rescaled such that the peak at angle 0 represents regions aligned with the flow.

#### Junction analysis in flow experiments

Cells were immunostained for Actin and VE-Cadherin and imaged on using a spinning disc confocal microscope as described above. Using a custom MATLAB script, cell junctions were manually annotated into 3 classes: 1) linear, 2) focal adherens, 3) reticular, based on VE-Cadherin morphologies as described in the main text. Having annotated junction morphologies, the fraction of the junction perimeter occupied by each type of junction was calculated. Cells whose boundaries did not fit in a frame or were difficult to annotate for several reasons (including weak/bad immunostaining, highly overlapping cells, dividing or dead cells) were excluded from analysis.

#### Aspect ratio of cells

Using cell boundaries, we also measured the aspect ratio of each cell (ratio of the major to the minor axis of cells), which depicts how elongated or circular a cell is. A higher aspect ratio value represents increased cell elongation. Furthermore, the mean intensities of actin and FHL2 per cell were measured as needed.

#### FHL2 intensity in cells

The mean FHL2 intensity in each cell was normalised to the respective actin intensity. For FHL2 on stress fibres, regions on fibres were highlighted and the intensity of FHL2 and actin were measured within the region (on a stress fibre). For western blotting, FHL2 signal was normalised to the loading control and the experimental mean of the respective controls.

#### k-means clustering

From measurements of cell junction fractions (linear, reticular, and focal adherens junctions) and cell aspect ratio, cells were clustered using a k-means algorithm on MATLAB using squared Euclidean distance, with k=2. These distinctly represent UF-like and DF-like clusters as they were matched with the actual experimental condition shown in Fig. S3B. Cells from all conditions are partitioned into two distinct clusters based on their similarities to one another and to the centroids. Distances of each point to the centroid were also depicted.

#### Linescan intensity profiles

For Fig. 2F, using background (100) subtracted intensities of FHL2 and actin, the FHL2 intensity was normalized to the actin intensity of each cell. For linescans in Fig. 2G, we measured the intensity profile of FHL2 and actin across a line drawn perpendicular to a few actomyosin stress fibres (∼3-5 fibres). The FHL2 and actin intensities were normalized such that the max intensity peak is set to 1. In the cells where line scans were drawn, FHL2 intensity in the entire cell was measured, and was around 6.5 times lower in the case of UF cell. Thus, to accurately represent the intensity variations of FHL2 between the conditions as well, FHL2 intensity in the UF condition was divided by 6.5. For Fig. 6B, GEF-H1 and actin intensities were measured across a line spanning few microtubules (∼4-6 microtubules). To account for variations in intensities across conditions, the GEF-H1 and tubulin intensities were normalized to the maximum intensity measured for each channel, i.e., the maximum intensity of GEF-H1 and tubulin were both set to 1.

#### Phosphorylated myosin light chain (pMLC) levels

The mean intensity of sparse cells and endothelial monolayers stained for pMLC Serine 19 (S19) residue were measured and normalized to the mean actin intensity of each cell to account for variability across experimental conditions. For western blotting, pMLC signal was normalised to the total MLC levels, and further normalised the experimental mean of the respective controls.

#### Traction force measurements

Traction forces and strain energies were analysed using a custom-designed macro in Fiji based on previous studies^80,105^. Z-stacks of beads before and after cell trypsinization were selected and aligned using the Align Slices in the Stack plugin in Fiji (a normalized cross-correlation algorithm). The bead displacements were computed from bead movements using particle image velocimetry (PIV). PIV analysis parameters were three interrogation windows of 128, 64 and 32 pixels with a correlation of 0.60. Using Fourier transform traction cytometry, a Young’s modulus of 8.4 kPa, a regularization factor of 10^−9^ and a Poisson ratio of 0.5, traction forces were calculated from the bead displacement fields.

#### GEF-H1 localisation

By drawing a small region on the edge of a cell protrusion where individual microtubules can be easily segmented, a binary mask was created using the tubulin channel. This mask was transferred on to the GEF-H1 channel, and the ratio of the intensity of GEF-H1 within the mask (on microtubules) and outside the mask (outside microtubules) was calculated. Regions of dense microtubules were excluded from analysis.

#### Microtubule correlation

Protrusion regions were marked in live-cell imaging movies of endothelial cells with spyTubulin. The tubulin correlation between each frame for the first 5 frames were calculated and averaged using a custom MATLAB script. To minimize variability across different experimental conditions and cell lines, the average correlation for each cell was normalised to the mean of the respective control cells in each experiment.

#### Microtubules at the cell edge

In control or FHL2 knockdown cells, a square region of a cell protrusion was marked, and microtubules were thresholded in the region. The area occupied by microtubules was measured.

