## Supplemental Information for "Mechanosensitive FHL2 tunes endothelial function"

**This PDF file includes:**
Figures S1 to S6 and corresponding figure legends
Supplemental Tables S1 and S2 and corresponding legends

#### Supplementary figure legends

**Figure S1: Actin- and adhesion- related differentially expressed genes in response to flow. (A)** Actin-related genes present within the top 10 BP, CC or MF most enriched actin-related GO terms which are significantly different between endothelial cells under UF or DF are shown. The size of the dot reflects the -log10(adjusted p-value). **(B)** Adhesion-related genes present within the top 10 BP, CC or MF most enriched adhesion-related GO terms which are significantly different between endothelial cells under UF or DF are shown. The size of the dot reflects the -log10(adjusted p-value). **(C)** Dotplot showing all genes encoding for LIM domain proteins which were detected in endothelial cells under DF or UF. Genes are ordered by adjusted p-value and logFC. The size of the dots indicates absolute logFC values. The left-most dot represents the most significantly upregulated gene in DF, and the right-most dot is the most significantly downregulated gene in DF encoding for a LIM domain protein (orange = padj < 0.05 & logFC > 0, purple = padj < 0.05 & logFC < 0, yellow = padj = > 0.05). Genes of the LIM family that are not expressed in TeloHAECs are also shown.

**Figure S2: FHL2 expression and localisation in endothelial cells. (A)** Western blot of lysates from TeloHAECs subjected to UF and DF for 24 h. Samples were analysed by immunoblotting for FHL2 and GAPDH (loading control). **(B)** Normalised FHL2 intensity in UF and DF cells. N=3 independent experiments. Dots represent experimental means, whiskers show the minimum and maximum values, and the line within represents the median. **(C)** Western blot of lysates from Ctl (-dox) and FHL2-overexpressing (+dox; FHL2 OE) cells. Samples were analysed by immunoblotting for FHL2 and tubulin (loading control). **(D)** Graph shows normalised FHL2 intensity in Ctl and FHL2 OE cells. N=3 independent experiments. Dots represent experimental means, whiskers show the minimum and maximum values, and the line within represents the median. **(E)** Immunostaining of FHL2 and actin, and visualisation of mEmerald-FHL2 tagging in Ctl and FHL2-OE cells. **(F)** Ctl and FHL2 OE cells were subjected to UF and immunostained for FHL2 and actin. Graph shows normalised FHL2 intensity of each cell under UF in both conditions. N=2 independent experiments (n=253 for Ctl and 236 for FHL2 OE). **Scale:** (E) 10 µm. **Statistical tests:** (B, D) Unpaired t-test. (E) Unpaired t-test followed by Mann-Whitney test. ****p<0.0001, **p<0.01, *p<0.05.

**Figure S3: Clustering analysis shows cell populations in UF-like and DF-like clusters. (A)** Using measurements of aspect ratio and fractions of the three junction types in Fig. 3B-D, cells were clustered using k-means, with k=2. Two clusters and the corresponding data points are shown. N>3 independent experiments (n=1179 cells clustered into Cluster 1 or Cluster 2). Graphs showing the normalised aspect ratio vs. the fraction of the cell perimeter occupied by focal adherens junctions. Each cell is represented as a dot, and all the WT cells under UF and DF from across various experiments were clustered and are represented on the graph. Cells are colour coded based on whether they were correctly categorised into Clusters 1 or 2, based on the experimental condition. All the grey dots show that the cells that were under UF or DF were clustered into UF-like Cluster 1 or DF-like Cluster 2 respectively. The pink dots represent a mismatch in the clustering where the cells are clustered different from the actual experimental condition. **(B)** Graphs showing the aspect ratio (ratio of the major to the minor axis of a cell) vs. the fraction of the cell perimeter occupied by reticular junctions. N=2 independent experiments (n=253 for UF Ctl, 256 for UF FHL2 OE). **(C, D)** UF Ctl and FHL2 OE cells were clustered (unbiased) into the pre-defined UF-like Cluster 1 (C1) or DF-like Cluster 2 (C2). Graphs showing the normalised aspect ratio vs. the fraction of the cell perimeter occupied by focal adherens junctions in (C) Ctl, (D) FHL2 OE cells. (C) UF Ctl cells completely belong to UF-like Cluster 1. (D) FHL2 OE cells were clustered into the predefined clusters, and the circles represent the dominant cluster (in this case, DF-like Cluster 2; C2), while the ‘x’ depicts the minority cluster (in this case, UF-like Cluster 1; C1). **Statistical tests:** Unpaired t-test. ns: p value not significant.

**Figure S4: Knockdown of FHL2 partially reverses athero-prone DF phenotypes. (A)** Western blot of lysates from control (Ctl; -dox) and FHL2-knockdown (shFHL2; +dox) TeloHAECs. Samples were analysed by immunoblotting for FHL2 and tubulin (loading control). **(B)** Normalised FHL2 intensity in Ctl and shFHL2 cells. N=4 independent experiments. Dots represent experimental means, whiskers show the minimum and maximum values, and the line within represents the median. **(C)** Inverted contrast images of Ctl and shFHL2 cells immunostained for actin and FHL2. **(D)** Fluorescence intensity of FHL2 normalised to actin intensity of each cell, in Ctl and shFHL2 TeloHAECs. N=5 independent experiments (n=187 for Ctl and 151 for shFHL2). **(E)** Graph shows the aspect ratio (ratio of the major to the minor axis of a cell) of control and shFHL2 (+dox) cells under DF. N=2 independent experiments (n=171 for Ctl, 225 for shFHL2). **(F)** From brightfield images, the alignment of Ctl and shFHL2 cells under DF are calculated by local gradient orientation, and represented as the probability distribution of cells with the flow alignment angles (centred around $0^o$). Shaded error bars representing standard deviation are shown. N=1 independent experiment (n=32 for UF Ctl, and n=39 for shFHL2; fields of view, with >100 cells/field). **(G)** Inverted contrast confocal images of control (Ctl; -dox) and shFHL2 TeloHAECs under DF, stained for phalloidin and VE-Cadherin. Annotated junctions are depicted as in Fig. 3A (purple: linear junctions, yellow: focal adherens junctions, orange: reticular junctions). **(H, I, J)** Graphs showing the aspect ratio (ratio of the major to the minor axis of a cell) vs. the fraction of the cell perimeter occupied by linear junctions (H), focal adherens junctions (I), reticular junctions (J). N=2 independent experiments (n=171 for DF Ctl, 225 for DF shFHL2 OE). The DF Ctl means were pooled from 5 experiments for statistical tests. **(K, L)** DF Ctl and shFHL2 cells were clustered (unbiased) into the pre-defined UF-like Cluster 1 (C1) or DF-like Cluster 2 (C2). Graphs showing the normalised aspect ratio vs. the fraction of the cell perimeter occupied by focal adherens junctions in (K) Ctl, (L) shFHL2 cells. (K) DF Ctl cells almost completely belong to DF-like Cluster 2 (circles represent the dominant cluster, in this case, DF-like Cluster 2, C2; while the ‘x’ depicts the minority cluster (in this case, UF-like Cluster 1; C1). (L) DF shFHL2 cells were clustered into the predefined clusters, and the circles represent the dominant cluster (in this case, UF-like Cluster 1; C1), while the ‘x’ depicts the minority cluster (in this case, DF-like Cluster 2; C2). **Scale:** (A, G) 10 µm. **(G) Statistical tests:** (B) Unpaired t-test. (D) Paired, two-tailed t-test. (E, H, I, J) Unpaired t-test followed by Mann-Whitney test. (F) Kolmogorov-Smirnov test. *p<0.05, **p<0.01, ***p<0.001, ns: p value not significant.

**Figure S5: FHL2 mutant and clustering analysis shows its similarity with UF-like Cluster 1. (A)** Inverted contrast images of FHL2-overexpressing (FHL2 OE; +dox) and FHL2-F1-4A-overexpressing (F1-4A OE; +dox) cells immunostained for VE-Cadherin. mEmerald tagging of FHL2 protein is visualised. **(B)** Western blot of lysates from control (Ctl; -dox) and F1-4A OE (+dox) cells. Samples were analysed by immunoblotting for FHL2 and tubulin (loading control). N=2 independent experiments. **(C)** Graph showing the aspect ratio (ratio of the major to the minor axis of a cell) vs. the fraction of the cell perimeter occupied by reticular junctions. N=2 independent experiments (n=143 for UF Ctl, 167 for UF F1-4A OE). One data point belonging to UF Ctl (aspect ratio 18.25, linear 0.62, focal adherens 0.01) used in the analyses is excluded from the graph for representation purposes. The UF Ctl means were pooled from 5 experiments for statistical tests. **(K, L)** UF Ctl and UF F1-4A OE cells were clustered (unbiased) into the pre-defined UF-like Cluster 1 (C1) or DF-like Cluster 2 (C2). Graphs showing the normalised aspect ratio vs. the fraction of the cell perimeter occupied by focal adherens junctions in (K) Ctl, (L) F1-4A OE cells. UF Ctl and UF F1-4A OE cells almost completely belong to UF-like Cluster 1 (circles represent the dominant cluster, in this case, UF-like Cluster 1, C1; while the ‘x’ depicts the minority cluster (in this case, DF-like Cluster 2; C2). **Scale:** (A) 20 μm. **Statistical tests:** (C) Unpaired t-test. ns: p value not significant.

**Figure S6: FHL2 and pMLC levels upon ROCK inhibition using Y-27632.** **(A)** Western blot of lysates from Ctl and shFHL2 cells, with or without Rho-kinase inhibitor (Y-27632 a.k.a. Y-27) are shown. Samples were analysed by immunoblotting for FHL2, pMLC, MLC, and tubulin (loading control). **(B)** Normalised pMLC intensity (pMLC/MLC intensity) in Ctl and shFHL2 cells treated with DMSO or Y-27. N=1 independent experiment.

#### Supplementary tables

**Table 1: Supplementary Table GO results.** Table shows the test results for enrichment of differentially expressed genes (adjusted p-value < 0.05) between endothelial cells under UF and DF in gene ontology (GO) terms using clusterProlifer in R software. p-value = result of Fisher’s exact test, qvalue = Benjamini-Hochberg corrected p-value to (multiple hypothesis test correction), geneID = names of DEGs annotated to the term, Count = number of DEGs in term. k.K. = number of DEG genes in GO term/number of genes in the GO term

**Table 2: Supplementary Table DEG results DESeq2.** Table shows the results of the DESeq2 differential gene expression test between endothelial cells under UF and DF. baseMean = mean of normalized counts for all samples, lfcSE = standard error, stat = Wald test statistic, p-value = p-value of the Wald test, padj = BH adjusted p-values.

**Figure S1**

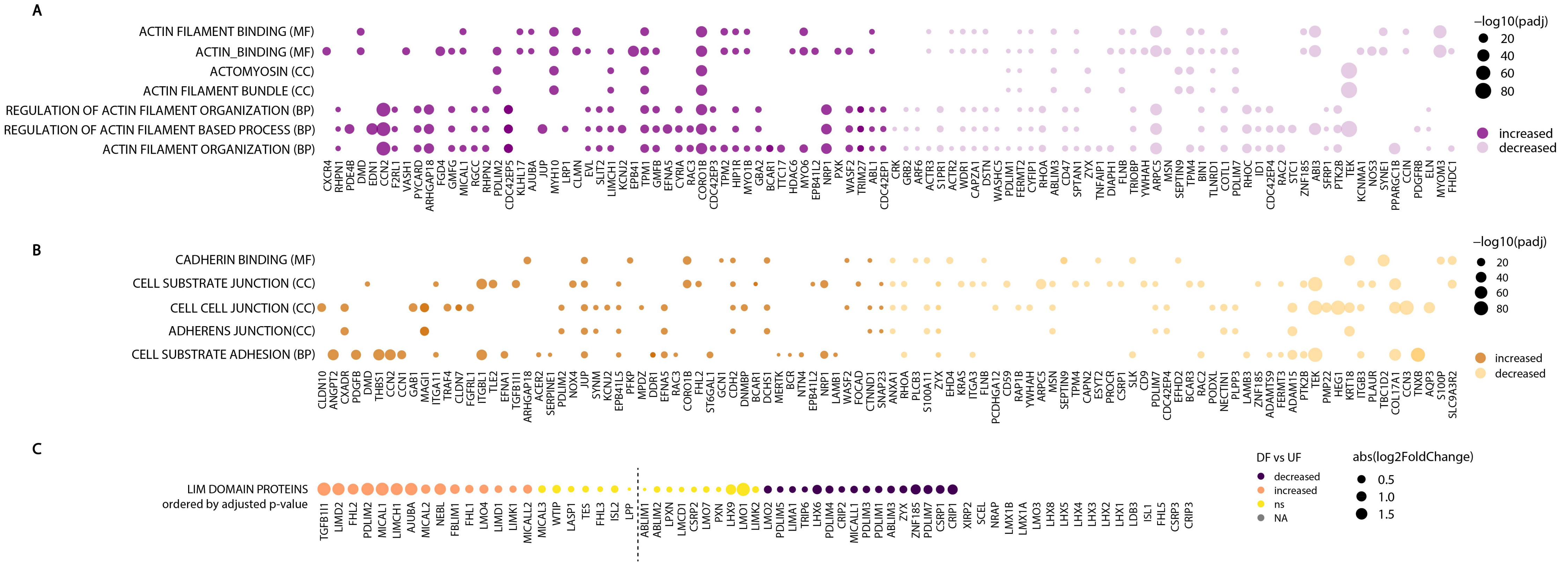
A
ACTIN FILAMENT BINDING (MF)
ACTIN_BINDING (MF)
ACTOMYOSIN (CC)
ACTIN FILAMENT BUNDLE (CC)
REGULATION OF ACTIN FILAMENT ORGANIZATION (BP)
REGULATION OF ACTIN FILAMENT BASED PROCESS (BP)
ACTIN FILAMENT ORGANIZATION (BP)
−log10(padj)
20
40
60
80
increased
decreased
B
CADHERIN BINDING (MF)
CELL SUBSTRATE JUNCTION (CC)
CELL CELL JUNCTION (CC)
ADHERENS JUNCTION(CC)
CELL SUBSTRATE ADHESION (BP)
−log10(padj)
20
40
60
80
increased
decreased
C
LIM DOMAIN PROTEINS
ordered by adjusted p-value
DF vs UF
decreased
increased
ns
NA
abs(log2FoldChange)
0.5
1.0
1.5

### Figure S2

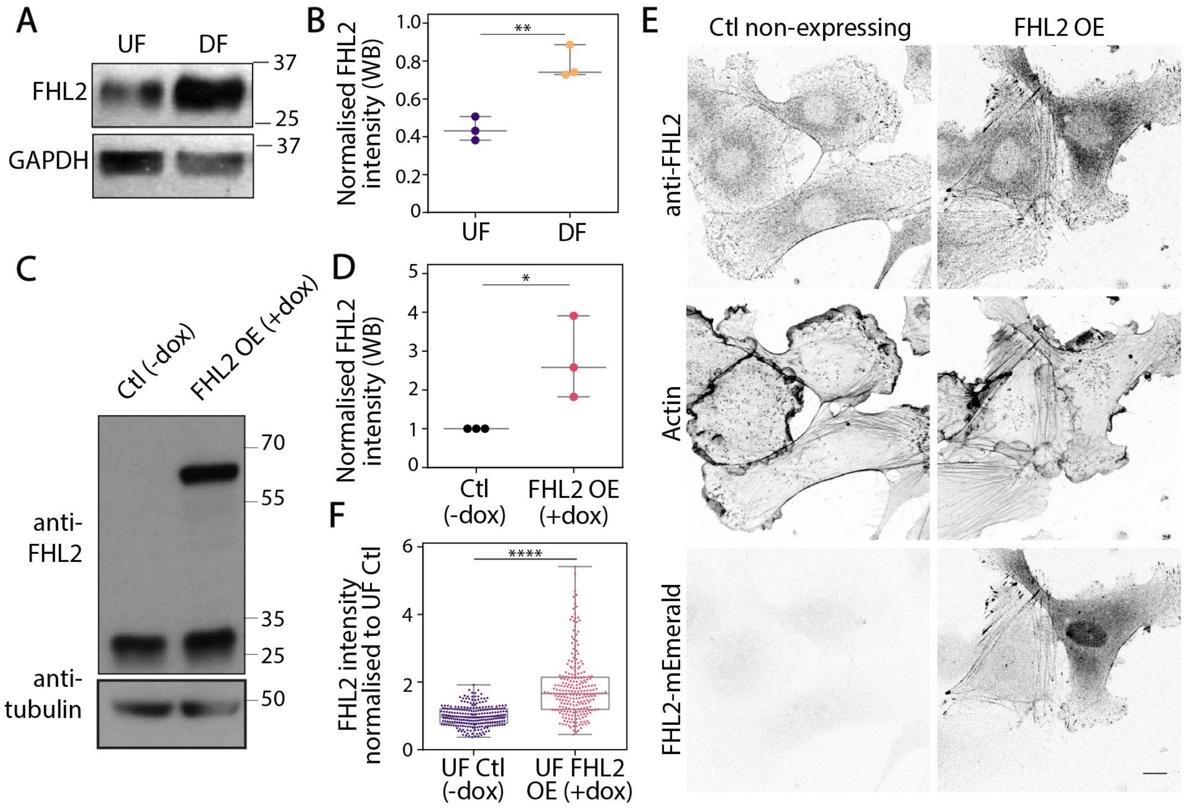

A
UF
DF
FHL2
GAPDH
37
25
37
B
Normalised FHL2 intensity (WB)
**
UF
DF
C
Ctl (-dox)
FHL2 OE (+dox)
anti-FHL2
anti-tubulin
70
55
35
25
50
D
Normalised FHL2 intensity (WB)
*
Ctl (-dox)
FHL2 OE (+dox)
F
FHL2 intensity normalised to UF Ctl
****
UF Ctl (-dox)
UF FHL2 OE (+dox)
E
Ctl non-expressing
FHL2 OE
anti-FHL2
Actin
FHL2-mEmerald

### Figure S3

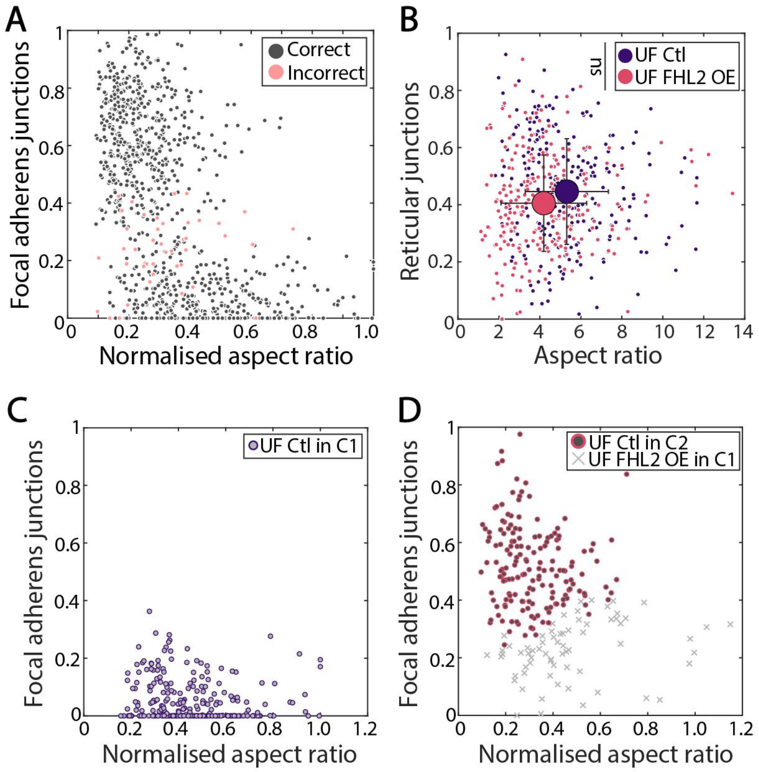
A
Focal adherens junctions
Normalised aspect ratio
Correct
Incorrect
B
Reticular junctions
Aspect ratio
ns
UF Ctl
UF FHL2 OE
C
Focal adherens junctions
Normalised aspect ratio
UF Ctl in C1
D
Focal adherens junctions
Normalised aspect ratio
UF Ctl in C2
UF FHL2 OE in C1

### Figure S4

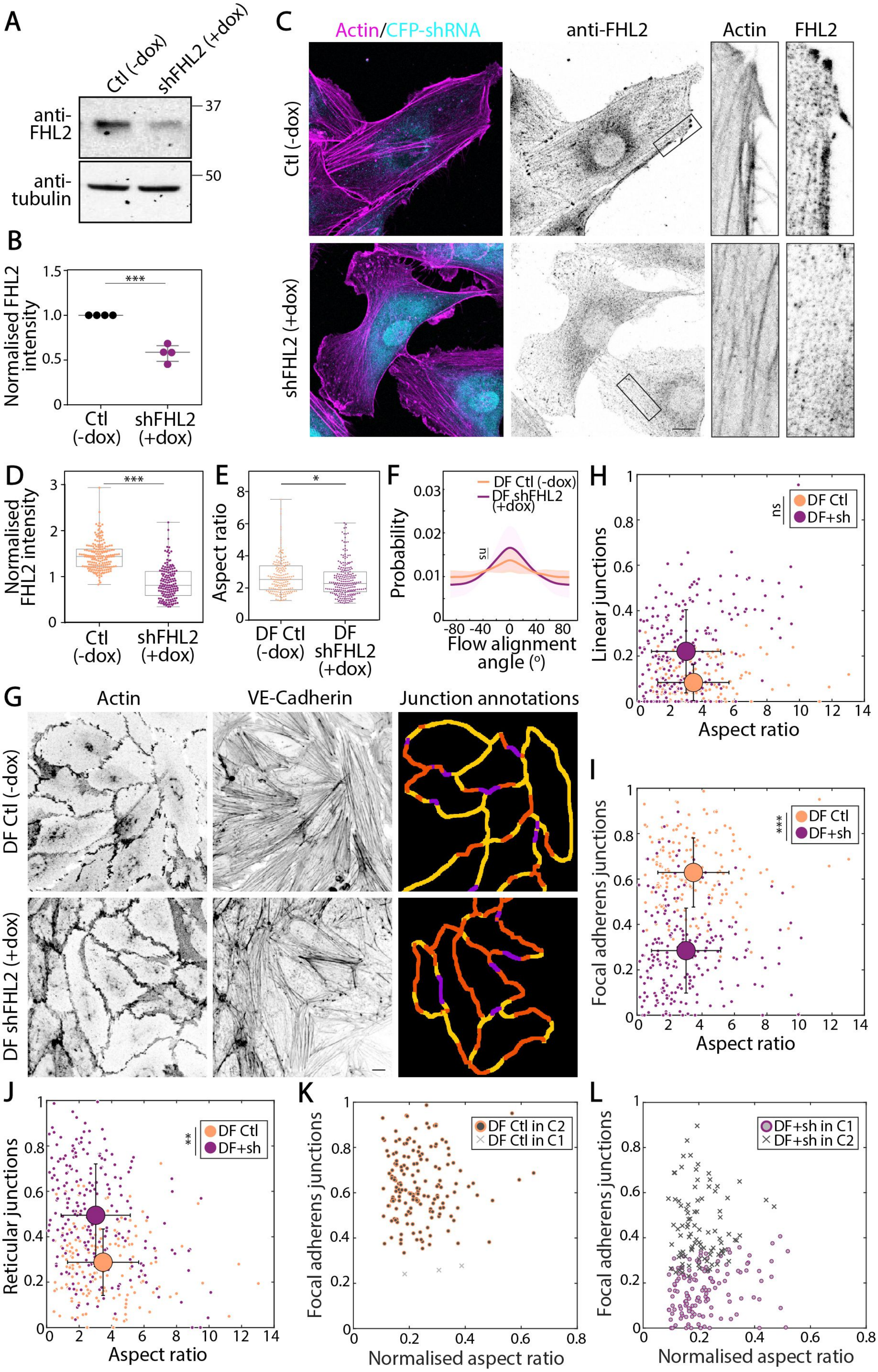
A
Ctl (-dox)
shFHL2 (+dox)
anti-FHL2
anti-tubulin
37
50
B
Normalised FHL2 intensity
***
Ctl (-dox)
shFHL2 (+dox)
C
Actin/CFP-shRNA
anti-FHL2
Actin
FHL2
Ctl (-dox)
shFHL2 (+dox)
D
Normalised FHL2 intensity
***
E
Aspect ratio
*
DF Ctl (-dox)
DF shFHL2 (+dox)
F
DF Ctl (-dox)
DF shFHL2 (+dox)
Probability
ns
Flow alignment angle (°)
G
Actin
VE-Cadherin
Junction annotations
DF Ctl (-dox)
DF shFHL2 (+dox)
H
Linear junctions
ns
DF Ctl
DF+sh
Aspect ratio
I
Focal adherens junctions
***
DF Ctl
DF+sh
Aspect ratio
J
Reticular junctions
**
DF Ctl
DF+sh
Aspect ratio
K
Focal adherens junctions
DF Ctl in C2
DF Ctl in C1
Normalised aspect ratio
L
Focal adherens junctions
DF+sh in C1
DF+sh in C2
Normalised aspect ratio

### Figure S5

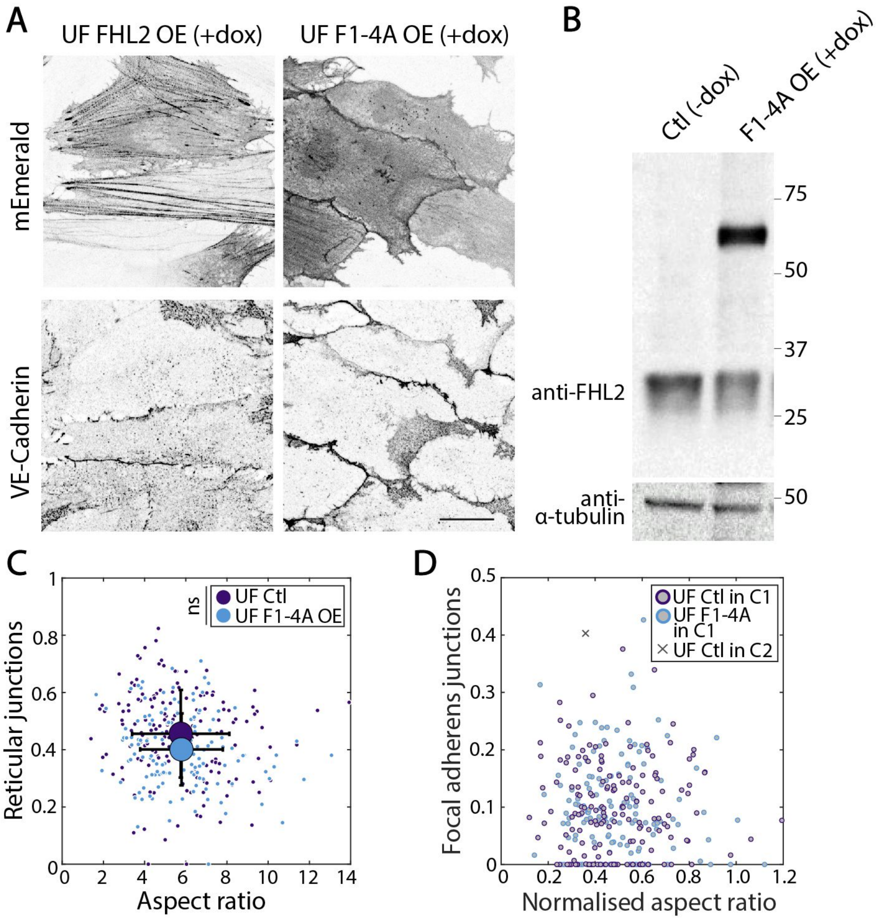
A
UF FHL2 OE (+dox)
UF F1-4A OE (+dox)
mEmerald
VE-Cadherin
B
Ctl (-dox)
F1-4A OE (+dox)
75
50
37
25
anti-FHL2
anti-
α-tubulin
50
C
ns
UF Ctl
UF F1-4A OE
Reticular junctions
Aspect ratio
D
UF Ctl in C1
UF F1-4A
in C1
UF Ctl in C2
Focal adherens junctions
Normalised aspect ratio

### Figure S6

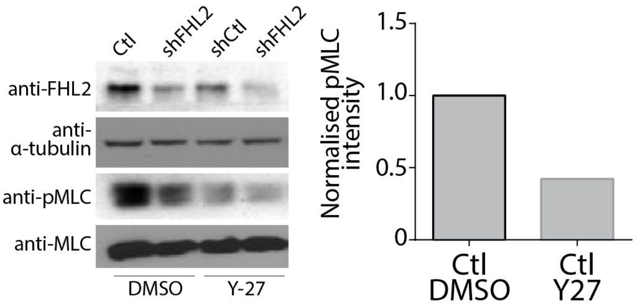

Ctl
shFHL2
shCtl
shFHL2
anti-FHL2
anti-
α-tubulin
anti-pMLC
anti-MLC
DMSO
Y-27
Normalised pMLC intensity
1.5
1.0
0.5
0
Ctl DMSO
Ctl Y27
